# Potential multiple disease progression pathways in female patients with Alzheimer’s disease inferred from transcriptome and epigenome data of the dorsolateral prefrontal cortex

**DOI:** 10.1101/2024.03.24.586514

**Authors:** Kousei Honda, Akinori Awazu

**Author notes:** Send correspondence to: Akinori Awazu.

## Abstract

Late-onset Alzheimer’s disease (AD) is a typical type of dementia for which therapeutic strategies have not yet been established. The database of the Rush Alzheimer’s Disease study by the ENCODE consortium contains transcriptome and various epigenome data. Although the Rush AD database may contain a satisfactory amount of data for women, the amount of data for men remains insufficient. Here, based on an analysis of publicly available data from female patients, this study found that AD pathology appears to be nonuniform; several different typical forms of cognitive deterioration were observed. First, cluster analysis was performed on individuals diagnosed with “No Cognitive Impairment (NCI),” “Mild Cognitive Impairment (MCI),” and “Alzheimer’s Disease (AD)” stages in clinical trials using gene expression. NCI was found to contain at least two substages, whereas MCI and AD contained four and six substages, respectively. The epigenome data, in particular genome-wide H3k4me3 distribution data, also supported the existence of multiple AD substages. However, *APOE* gene polymorphisms of individuals seemed to not correlate with disease stage. An inference of adjacency networks among substages, evaluated via partition-based graph abstraction using the gene expression profiles of individuals, suggested the possibility of multiple typical disease progression pathways from NCI to different AD substages through various MCI substages. These results will aid future research on the pathological features and mechanisms of AD and may induce changes in the strategic bases of conventional AD prevention and treatment.

## INTRODUCTION

Alzheimer’s disease (AD) is the most common cause of dementia, accounting for 60– 70 % of dementia cases in older adults [1, 2]. AD can be classified into the familial type, which is strongly influenced by genetic factors, and the late-onset (sporadic) type, which is strongly influenced by aging and environmental factors, with the latter accounting for the majority of AD cases [3]. No treatment or preventive methods for AD have yet been established because the fundamental pathogenic mechanism of sporadic AD remains unclear.

Plaques of abnormally aggregated amyloid β (Aβ) are often observed in the brains of patients with AD and are considered to be related to the onset and progression of the disease [4-7]. However, cases have been reported in which AD developed without the accumulation of Aβ [8] and inhibiting Aβ production even worsened symptoms (Francesca et al., 2023). Factors other than Aβ plaque formation, such as the abnormal aggregation of tau protein [9], the expression of apolipoprotein E variant APOE4 [10], herpes virus and coronaviruses (including SARS-CoV2) infections [11-14], diabetes [15, 16], brain inflammation [17], oxidative damage in cells [18-20], endoplasmic reticulum and ciliary dysplasia [21-24], myelin dysplasia [25, 26], and retrotransposons activities [27-29] have also been reported to influence the onset and progression of AD. Owing to the involvement of the aforementioned factors, the causes and symptoms of AD are thought to vary widely from patient to patient. Further, variations have been previously reported in symptoms and disease progression rates among patients [30-33].

In recent years, transcriptome and genome-wide analysis data of nucleotide polymorphisms have been collected from patients with AD, and various differentially expressed genes and patient-specific SNPs have been identified in healthy individuals and patients with AD [34-36]. Notably, these characteristics are not always seen equally across all studies. One reason for this is the variations in technical factors among studies. However, the diversity in molecular conditions and gene expression patterns in the brain among patients with AD could also be a major contributor [37-41].

Thus, AD progression is expected to exhibit various disease states. Classifying and characterizing these disease states and progression pathways is necessary to enable appropriate state-dependent treatment. In addition to a comparison of the features of the entire transcriptome, a comparison of the entire epigenome status among patients should be useful for such classification and characterization of disease diversity. The database of the Rush Alzheimer’s Disease study by the ENCODE consortium (https://www.encodeproject.org/) [42, 43] contains transcriptome and various epigenome data such as histone modifications (H3K27ac, H3K27me3, and H3K4me3) and CCCTC-binding factor (CTCF) binding site distribution from individuals in the “No Cognitive Impairment (NCI),” “Mild Cognitive Impairment (MCI),” and “Alzheimer’s Disease (AD)” groups that are available for unlimited public use (termed Rush AD database thereafter). Therefore, analyzing these publicly available data of patients with different cognitive impairment stages is expected to elucidate the diversity in the onset and progression of late-onset AD in more detail.

However, although the Rush AD database seemed to contain a satisfactory amount of data for women, the amount of data for men seemed insufficient, particularly for patients with MCI and AD, where the number of female patients in NCI, MCI, and AD groups was 25, 29, and 28, respectively, whereas that of male patients was 14, 4, and 7, respectively. Therefore, in this study, the classification of disease status and AD progression pathways were performed using transcriptome and epigenome data of female patients in the Rush AD database as a first step of the analysis of the diversity of patients with AD using multiomics data. Our findings are expected to be useful not only for female patients but also for understanding symptoms in male patients, even though some features such as disease progression rate are different between male and female patients [44, 45].

In this study, patient diversity was considered by assessing the deviation from healthy individuals and diversity of disease progression. For this purpose, we first searched for differentially expressed gene sets between healthy individuals and patients. The gene sets searched here were those which contained the largest number of genes from gene sets that could actually distinguish healthy individuals from patients using cluster analysis. The candidate genes included therein were selected by identifying statistically significant differences in their expression levels. Second, we focused on the variety in the expression patterns of those gene sets between patients to show the existence of diverse conditions and subgroups within patients. The existence of these subgroups was also supported by comparisons of epigenomic data between individuals. Third, the relationship between subgroups and structure of the disease progression pathway was predicted based on the similarity in gene expression between subgroups.

## RESULTS

### Fundamental features of the analyzed transcriptome and epigenome data

RNA sequencing (RNA-seq) data (read count and TPM) of the dorsolateral prefrontal cortex of female individuals belonging to the NCI, MCI, and AD groups were acquired from the Rush AD database. In this dataset, the number of RNA-seq data of individuals in the NCI, MCI, and AD groups were 25, 29, and 28, respectively (Table 1, Table S1a– S1b).

**Table 1.** Summary of the individuals included in this study (see Table S1a and S1b for more details).

| Clinical stage | No Cognitive Impairment (NCI) | Mild Cognitive Impairment (MCI) | Alzheimer’s Disease (AD) |
| --- | --- | --- | --- |
| # of individuals | 25 | 29 | 28 |
| Individual index | N1 – N25 | M1 – M29 | A1 – A28 |

To examine the quality and bias of analyzed data, we estimated one of the size factors of the RNA-seq data, that is, the median of the ratios (MOR) defined in the recent literature for DESeq [46], using read count data (Table S1a). The ratio [standard deviation of MOR over individuals]/[average MOR over individuals] was 0.1075. Similarly, we evaluated the first and third quartiles of the ratio (1OR and 3OR) between the average and standard deviation, which were 0.1147 and 0.1006, respectively. These results suggested that the size factor would not considerably input bias among samples or affect the comparison of either the read count data or TPM values.

When examining the significance of the difference in gene expression levels between individuals aged <85 years and those >90 years, we did not detect any genes exhibiting a false discovery rate (FDR) < 0.1 (likelihood ratio test and two-tailed *t*-test) (Table S1a-S1b). This suggested that the variations in gene expression levels among the NCI, MCI, and AD groups was not due to the influence of age but rather due to differences in clinical stages.

In addition to RNA-seq data, the bigwig format data of fold change over control (FC) on genome-wide histone modifications (H3K4me3, H3K27ac, and H3K27me3) and CTCF binding site distribution obtained through chromatin immunoprecipitation sequencing (ChIP-seq) were acquired from the Rush AD database for 39 individuals: 11, 11, and 17 individuals from the NCI, MCI, and AD groups, respectively (Table S1c). In this study, the regions with FC > 5 were defined as the regions of peak intensities of these epigenome marks [47].

### Cluster analysis of differentially expressed gene sets in the majority of patients with MCI

To extract a differentially expressed set of genes among patients with NCI, MCI, or AD, we performed the likelihood ratio test, which is one of the most familiar tests for estimating differentially expressed genes (DEGs) from read count data, for each gene using the DESeq2 software [48] (Table S1a). Consequently, we performed hierarchical clustering of individuals using genes with P-values below the threshold value P^D^_NMA_ = 0.01, 0.02, 0.03, 0.04, and 0.05 (Figs. 1 and S1a–S1e). When P^D^_NMA_ ≤ 0.05, a cluster with significantly more patients with MCI (MCI cluster) and a cluster with patients with NCI, AD, and a small number of patients with MCI was formed via the first divergence of the dendrogram (P < 0.05, Fisher’s exact test; Fig. S1a–S1e). However, even though the gene set for clustering contained only genes with smaller P values, patients with NCI, MCI, and AD were not from their respective cluster; no clusters comprising solely of patients with NCI, MCI, or AD were formed through the first divergence of the dendrogram.

**Fig. 1.**
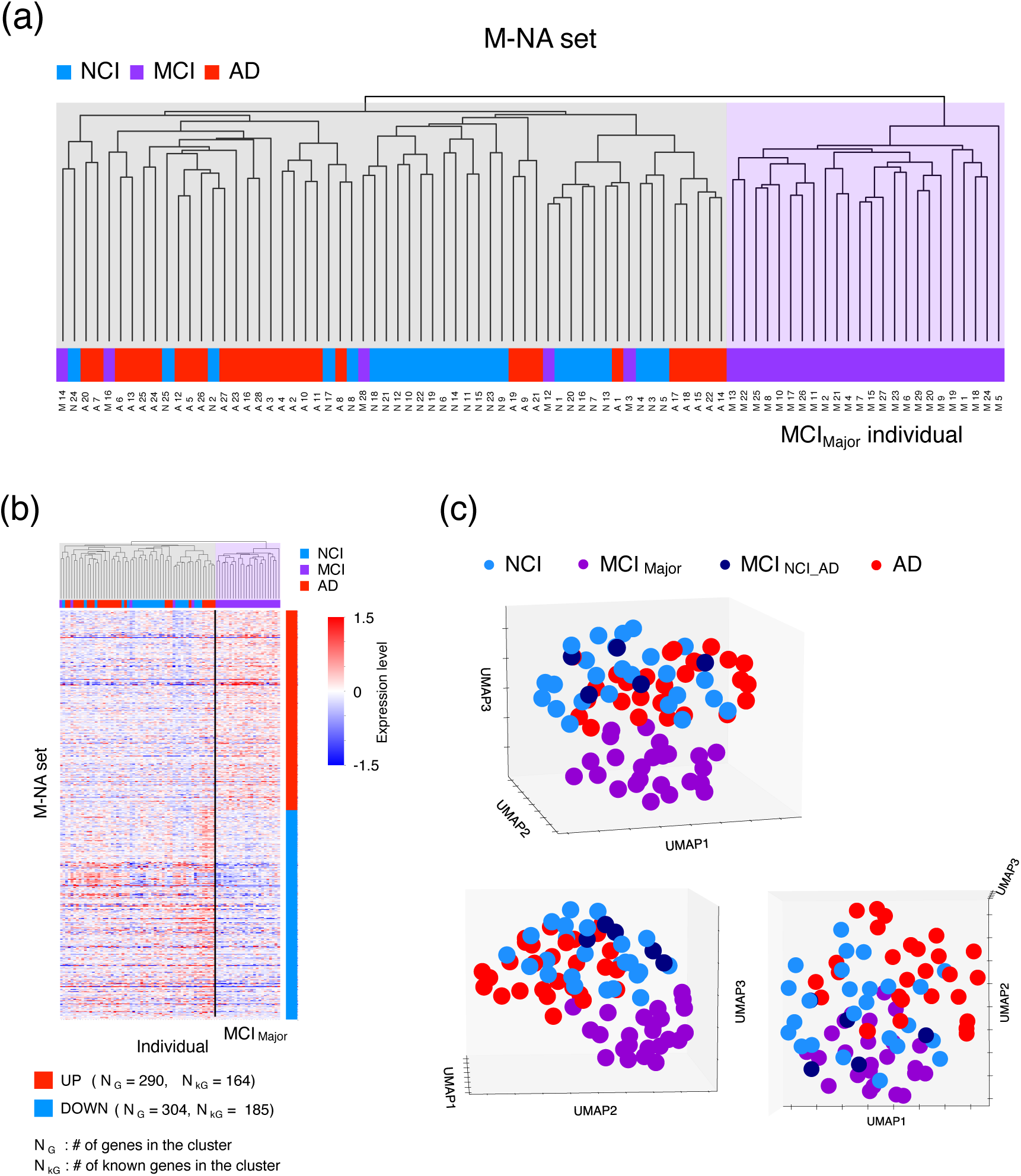
Results of the hierarchical clustering of individuals and of UMAP obtained using the M-NA set. Result of the hierarchical clustering of individuals when using the M-NA set (P^A^_NMA_ = 0.04) (a). The expression level of each gene in the M-NA set for each individual was plotted (b) (see Table S2a for more details). The heatmap of the expression levels of each gene was described using X = log_2_(TPM + 1) – C, where C was chosen to satisfy the average of X over individuals = 0 for each gene. MCI_Major_ individuals occupy the right region in the dendrogram. N_G_ and N_kG_ indicate the total number of genes and number of genes annotated as protein coding or noncoding RNA genes with higher (UP) and lower (DOWN) expression levels in MCI_Major_ individuals, respectively. Scatter plot of individuals in 3-dimensional space constructed using U-MAP with 1, 2, and 3 axes shown from various directions (c).

We then performed the same analysis using analysis of variance (ANOVA), which is the most familiar test for evaluating differences of characteristic values among three or more groups, to determine the P-values using TPM values (Table S1b), where P^A^_NMA_ was defined instead of P^D^_NMA_. Herein, ANOVA was generally better for detecting differences and was expected to identify more candidate genes in a gene set than the likelihood ratio test. Consequently, an MCI cluster and a cluster with patients with NCI, AD, and a small number of patients with MCI were formed, as in the previous case (Fig. S1f-S1j). Additionally, when the P^A^_NMA_ was 0.04, a cluster comprising solely patients with MCI was formed through the first divergence of the dendrogram (Fig. 1a). In contrast, we did not observe any clear separation between patients with NCI and those with AD, regardless of the P^A^_NMA_ value.

Notably, we did not obtain any gene set when applying the P^D^_NMA_ and P^A^_NMA_ values that enabled the complete discrimination of patients with NCI, MCI, and AD. In addition, depending on the P^D^_NMA_ and P^A^_NMA_ values, the patients with MCI who deviated from the MCI cluster varied; however, 24 patients with MCI were always included in the MCI cluster (Fig. S1). This suggests that the MCI cluster consisted of 24 patients with MCI, named MCI_Major_ individuals, whereas the 5 other patients with MCI (M3, M12, M14, M16, and M28) who tended to deviate from the MCI_Major_ individuals, were named MCI_NCI_AD_ individuals. We assumed that the MCI_Major_ and MCI_NCI_AD_ individuals were classified to the substages named MCI_Major_ and MCI_NCI_AD_, respectively.

Moreover, the set of genes with P-values of the above ANOVA ≤ P^A^_NMA_ = 0.04 was named the “M-NA set” as it significantly differed between the majority of patients with MCI from those with NCI and AD (Fig. 1b, Table S2a). In a 3-dimensional U-map using the M-NA set, we clearly observed the deviation of the MCI_Major_ individuals from those at other substages, with MCI_NCI_AD_ detected in a mixed group of patients with NCI and AD (Fig. 1c).

The classification result of patients with MCI into MCI_Major_ and MCI_NCI_AD_ individuals seemed to be independent of the age of individuals. For example, the numbers of patients over and under 90 years old in MCI_Major_ were 19 and 5, whereas those in MCI_NCI_AD_ were 3 and 2, respectively (P-value of the Fisher’s exact test = 0.5688; Table S1a–S1b).

### Functional features of genes in the M-NA set

We also considered the functional features of genes in the M-NA set. In this set, we obtained 594 genes (290 upregulated and 304 downregulated genes in MCI_Major_); of them, 349 genes (164 upregulated and 185 downregulated in MCI_Major_) were annotated (Fig. 1b) as protein coding or noncoding RNA genes using Entrez [49]. Herein, both sets of upregulated and downregulated genes in MCI_Major_ contained some previously reported AD risk-associated genes; in particular, the upregulated gene set contained *CYP19A1, GSTM1, HSD11B1, IL10, MT-RNR1, PCK1, TRAK2, UBE2D1*, and *XBP1*, whereas the downregulated gene set contained *MT-ATP8* and *NME8* [50].

In addition, we performed enrichment analysis of all genes including genes with higher and lower expression levels in MCI_Major_ individuals (Table S2b–S2d). We identified that genes with higher expression levels in MCI_Major_ individuals were enriched in GO terms for the functions “response to lipopolysaccharide,” “response to molecule of bacterial origin,” “Toll-like receptor signaling pathway,” “regulation of MHC class II biosynthetic process,” “vitamin metabolic process,” “water-soluble vitamin metabolic process,” “chemotaxis,” and “taxis” (P-value < 10^-3^ and FDR < 0.5) (Table S2c). Whereas, we identified that genes with lower expression levels in MCI_Major_ individuals were enriched in GO terms for the functions “mRNA 5′-splice site recognition” and “mRNA cis splicing, via spliceosome” (P-value < 10^-3^ and FDR < 0.5) (Table S2d). Recent studies have suggested that some of these functional features are characteristic of AD pathology; for example, the lipopolysaccharide found in the wall of all Gram-negative bacteria was reported to play a role in causing AD [51, 52], Toll-like receptors were found to participate in neuronal cell death by apoptosis [53, 54], MHC class II-expressing cells were suggested to be involved in the degradation of neurons [55], vitamins were reported to affect the generation and clearance of Aβ [56], and microglia were shown to exhibit chemotaxis for Aβ clearance [57, 58].

### Differentially expressed gene sets between patients with NCI and AD

To extract a differentially expressed gene set between patients with NCI and those with AD, we performed hierarchical clustering of individuals using the genes satisfying the following criteria: mean expression level (TPM) was higher in patients with AD than in those with NCI, with smaller P-values in the likelihood ratio test than the P^D^_NA_ threshold, or mean expression level (TPM) was lower in patients with AD than in those with NCI, with smaller P-values in the likelihood ratio test than the P^D^_AN_ threshold. Subsequently, we searched the sets of P^D^_NA_ and P^D^_AN_ values that divided the patients with NCI and those with AD into different clusters in the first branch of the dendrogram. Within the range P^D^_NA_ ≤ 0.05 and P^D^_AN_ ≤ 0.05, we identified the finite but narrow area of P^D^_NA_ and P^D^_AN_, P^D^_NA_ = 0.01 and P^D^_AN_ = 0.02, P^D^_NA_ = 0.01 and P^D^_AN_ = 0.03, P^D^_NA_ = 0.02 and P^D^_AN_ = 0.03, P^D^_NA_ = 0.03 and P^D^_AN_ = 0.03, and P^D^_NA_ = 0.05 and P^D^_AN_ = 0.02, where patients with NCI were completely separated from those with AD (Fig. 2a). Thus, we obtained the largest set of genes separating the NCI and AD groups at P^D^_NA_ = 0.05 and P^D^_AN_ = 0.02 (Fig. S2a, Table S3a); this gene set was named the N-A_D1_ set. Notably, the number of genes with a lower mean expression level (TPM) in patients with AD than in those with NCI was larger when P^D^_NA_ = 0.03 and P^D^_AN_ = 0.03 than when P^D^_NA_ = 0.05 and P^D^_AN_ = 0.02 (Fig. S2b, Table S3b). Notably, this gene set contained all genes in other gene sets with P^D^_NA_ and P^D^_AN_ smaller than 0.03 that could separate the NCI and AD groups. Thus, we focused on the gene set obtained at P^D^_NA_ = 0.03 and P^D^_AN_ = 0.03 and named it the “N-A_D2_ set.”

**Fig. 2.**
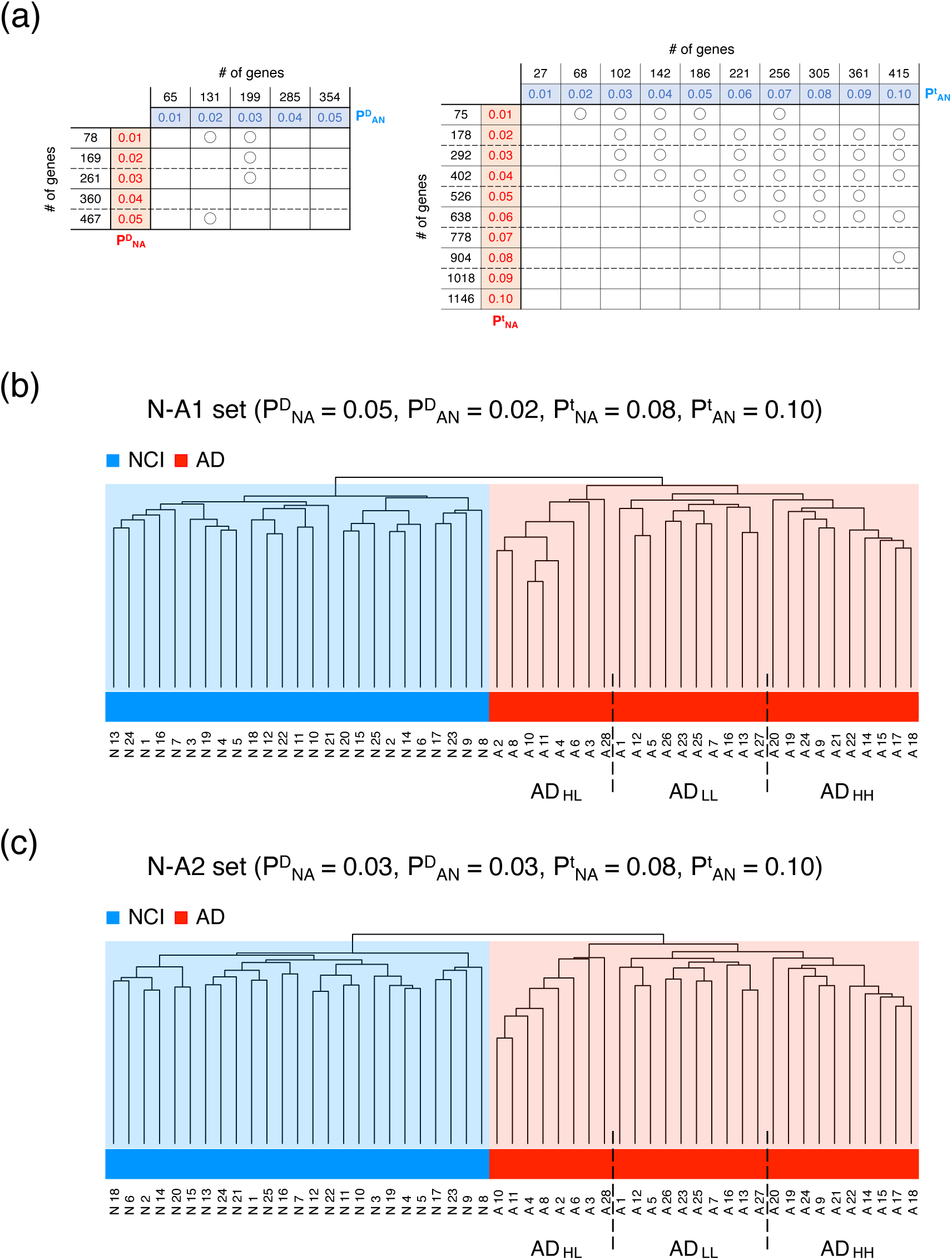
Phase diagrams of the hierarchical clustering of patients with NCI and AD, and the results of the hierarchical clustering of individuals in the N-A1 and N-A2 sets. Phase diagram of the results of hierarchical clustering when using genes where p ≤ P^D^_NA_ and p ≤ P^D^_AN_ (a, left), and that when using genes where p ≤ P^t^_NA_ and p ≤ P^t^_AN_ (a, right). Circles indicate the complete separation of patients with NCI from those with AD into different clusters at the first branch of the dendrogram. The dendrograms obtained using the N-A1 (b) and N-A2 sets (c) are shown. Broken lines indicate the boundary between the AD_HL_, AD_LL_, and AD_HH_ groups.

We performed the same analysis using two-tailed *t*-tests to determine the P-values, defining the P^t^_NA_ and P^t^_AN_ instead of the P^D^_NA_ and P^D^_AN_, respectively. Herein, the *t*-test was generally better for detecting differences and was expected to identify more candidate genes in a gene set than the likelihood ratio test. Searching in the ranges P^t^_NA_ ≤ 0.05 and P^t^_AN_ ≤ 0.05, patients with NCI were completely separated from those with AD in 12 of 15 cases of P^t^_NA_ and P^t^_AN_ set for P^t^_NA_ ≤ P^t^_AN_ (Fig. 2a). When we searched the wider range P^t^_NA_ ≤ 0.1 and P^t^_AN_ ≤ 0.1, patients with NCI were completely separated from those with AD in 37 of 55 cases of P^t^_NA_ and P^t^_AN_ set for P^t^_NA_ ≤ P^t^_AN_ (Fig. 2a). We thus obtained the largest set of genes separating the NCI and AD groups at P^t^_NA_ = 0.08, and P^t^_AN_ = 0.1 (Fig. S2c, Table S3c). This gene set was named the “N-A_t_ set.”

Hierarchical clustering using a combination of the expression levels of all genes in the N-A_t_ and N-A_D1_ sets also separated patients with NCI from those with AD (Fig. 2b, Table S3d). Conversely, the expression of all genes in the N-A_t_ and N-A_D2_ sets did not separate patients with NCI from those with AD. However, clustering using a combination of the expression levels of genes in the N-A_t_ set and a part of the N-A_D2_ set with low P-values completely separated patients with NCI from those with AD (Fig. 2c, Table S3e). Such gene sets are expected to involve various typical differentially expressed genes between patients with NCI and those with AD. In this study, the gene set consisting of the N-A_t_ and N-A_D1_ sets, which could completely separate patients with NCI from those with AD, was named the “N-A1 set” (Table S3d), whereas the gene set consisting of the N-A_t_ set and maximum number of genes in the N-A_D2_ set with low P-values was named the “N-A2 set” (Table S3e).

### Patients with AD can be classified into three groups

From the results of the hierarchical clustering for patients with NCI and AD performed using the N-A1 set, we noticed the emergence of three subgroups of patients with AD (Figs. 2b and 3a). Thus, we performed ANOVA on the expression levels of genes that were higher in patients with AD than in those with NCI among the three assumed subgroups (1020 genes). We identified 367 genes whose expression levels differed significantly (FDR < 0.1) among the 3 subgroups. Performing k-means cluster analysis of the expression levels of these genes in patients with AD, we identified two characteristic clusters: a cluster consisting of 96 genes the expression levels of which were low in only one subgroup (Cluster CA1) and another consisting of 271 genes the expression levels of which were high in only one subgroup (Cluster CB1) (Fig. 3a).

**Fig. 3.**
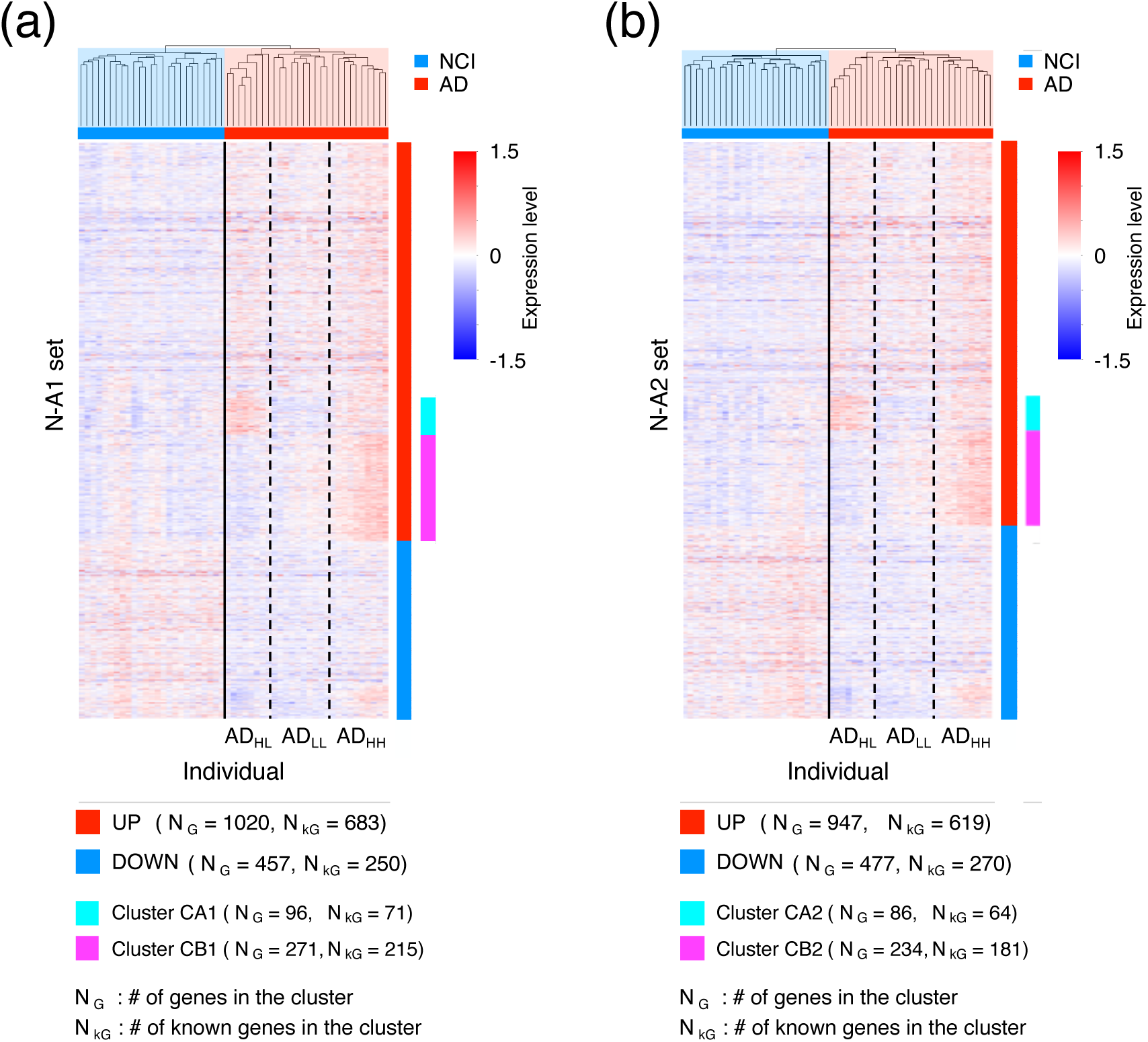
Results of the hierarchical clustering of individuals in the N-A1 and N-A2 sets and expression level of each gene. The dendrograms and expression level of each gene in the N-A1 set for each individual (a) (see Table S3d for more details) and those in the N-A2 set for each individual (b) (See Table S3e for more details) were plotted. The heatmap of the expression levels of each gene was described using X = log_2_(TPM + 1) – C, where C was chosen to satisfy the average of X over individuals = 0 for each gene. Broken lines indicate the boundary between the AD_HL_, AD_LL_, and AD_HH_ groups.

Thereafter, we performed a similar analysis for patients with NCI. Based on the shape of the dendrogram, patients with NCI were divided into three or four subgroups (Fig. 2b, Fig. 3a). However, when patients with NCI were divided into three subgroups, ANOVA revealed only nine genes with FDR < 0.1 among the three subgroups (Table S3d). Moreover, when patients with NCI were divided into four subgroups, ANOVA revealed only two genes with FDR < 0.1 among the four subgroups (Table S3d). Based on these values, the number of genes in clusters CA1 and CB1 was sufficiently large.

Collectively, these results suggested that the gene expression patterns in patients with AD, which can be classified into three types, were highly nonuniform. Patients with AD exhibiting high gene expression levels in both clusters, CA1 and CB1, were classified to the AD_HH_ subgroup (named AD_HH_ individuals), those exhibiting low gene expression levels in both CA1 and CB1 were classified in the AD_LL_ subgroup (named AD_LL_ individuals), whereas those exhibiting high gene expression levels in cluster CA1 and low gene expression levels in cluster CB1 were classified in the AD_HL_ subgroup (named AD_HL_ individuals).

Hierarchical clustering of patients with NCI and AD using the N-A2 set also revealed three possible subgroups (Figs. 2c and 3b). By performing the same analysis described above, we identified gene clusters similar to cluster CA1 containing 86 genes (cluster CA2) and cluster CB1 containing 234 genes (cluster CB2). Accordingly, patients with AD were divided into AD_HH,_ AD_LL_, and AD_HL_ groups based on the expression profiles of genes in clusters CA2 and CB2. The members of the AD_HH_, AD_LL_, and AD_HL_ groups were the same in both the N-A1 and N-A2 sets (Fig. S3a).

Importantly, variations were observed among the AD_HH_, AD_HL_, and AD_LL_ individuals when different P^t^_NA_ and P^t^_AN_ values were employed to determine the genes in the N-At set. However, for small changes in the P^t^_NA_ and P^t^_AN_ values, less variation in individuals was observed among subgroups; for example, when using P^t^_NA_ = 0.06 and P^t^_AN_ = 0.1 instead of P^t^_NA_ = 0.08 and P^t^_AN_ = 0.1 to determine the N-At. Herein, the gene set named N-A3 was defined as that consisting of genes in the N-A_t_ set and maximum number of genes in N-A_D1_ set with low P-values that could completely separate patients with NCI from those with AD, whereas the gene set named N-A4 was defined as that consisting of genes in the N-A_t_ set and maximum number of genes in N-A_D4_ set with low P-values that could completely separate patients with NCI from those with AD. The classification of patients with AD into the AD_HH_, AD_LL_, and AD_HL_ groups was similar to that obtained for N-A1, N-A2, N-A3, and N-A4, where 24 of the 28 individuals belonged to the same subgroups (Fig. S3). This suggested the robustness of our method in classifying the AD_HH_, AD_HL_, and AD_LL_ individuals.

In addition, the classification result of patients with AD into AD_HH_, AD_HL_, and AD_LL_ individuals seemed to be independent of the ages of individuals. For example, the numbers of individuals aged over and under 90 years in AD_HH_ were eight and two, those in AD_LL_ were six and four, and those in AD_HL_ were four and four, respectively (P-value = 0.4850).

### Common functional features of the N-A1 and N-A2 sets

We also considered the common functional features of the N-A1 and N-A2 sets. The N-A1 set contained 1477 genes (1020 upregulated and 457 downregulated, AD), with 933 genes (683 upregulated and 250 downregulated) annotated as protein coding or noncoding RNA genes using Entrez [49] (Fig. 3a). The N-A2 set contained 1424 genes (947 upregulated and 477 downregulated, AD), with 889 genes (619 upregulated and 270 downregulated) annotated as protein coding or noncoding RNA genes using Entrez [49] (Fig. 3b). The upregulated genes in both the N-A1 and N-A2 set contained *AR*, *CBS*, *CLU*, *CYP19A1*, *DBH*, *GNA11*, *HLA-A*, *IQCK*, *MPO*, *MS4A4E*, *MX1*, *NLRP3*, *SDC2*, *TAP2*, and *ZFHX3* (Table S3d–e), all of which have been reported to be AD risk-associated genes in recent studies [50, 59-61]. Similarly, the downregulated genes in both the N-A1 and N-A2 set contained *COL25A1* and *TARDBP* (Table S3d–e), which have also been reported to be AD risk-associated genes [50].

Additionally, the clusters CA1, CB1, CA2, and CB2 contained 71, 215, 64, and 181 annotated genes, respectively (Fig. 3). We identified *CLU*, *SDC2*, and *ZFHX3* in both the CA1 and CA2 clusters, whereas *DBH*, *GNA11*, *IQCK*, *MPO*, and *NLRP3* were identified in both the CB1 and CB2 clusters (Table S3d–e), thereby suggesting that these genes are responsible for the differences in the features among the three AD subgroups [50, 61].

Next, we performed enrichment analysis of all genes, including those with higher and lower expression levels in patients with AD for both the N-A1 and N-A2 set (Table S3f–S3o). We identified that genes with higher expression levels in patients with AD in both the N-A1 and N-A2 set were commonly enriched in GO terms for “forebrain dorsal/ventral pattern formation,” “mRNA 5′-splice site recognition,” and “negative regulation of NF-kappaB transcription factor activity” (P-value < 10^-3^ and FDR < 0.5) (Tables S3g and S3j). Whereas, genes with lower expression levels in patients with AD in both the N-A1 and N-A2 set were enriched in GO terms for “negative regulation of intrinsic apoptotic signaling pathway in response to DNA damage by p53 class mediator,” “negative regulation of DNA damage response, signal transduction by p53 class mediator,” and “coronary vasculature morphogenesis” (P-value < 10^-3^ and FDR < 0.5) (Table S3h, S3k). In the nervous system, NF-κB is known as one of the crucial components in the molecular switch that converts short- to long-term memory [62]. Thus, the upregulation of genes that suppress NF-κB expression is expected to exacerbate AD pathology. In addition, the downregulation of genes that suppress apoptosis in response to DNA damage would also promote AD progression through the enhancement of cell death in the central nervous system [63, 64].

We found that genes in both the CA1 and CA2 clusters were enriched in GO terms for “response to xenobiotic stimulus”, “cellular response to lipid”, “negative regulation of small molecule metabolic process”, “lung development”, “cellular response to external stimulus”, “negative regulation of carbohydrate metabolic process”, “neuron migration”, and “positive regulation of epithelial cell differentiation” (P-value < 10^-3^ and FDR < 0.5) (Tables S3l and S3n). Whereas, genes in both the CB1 and CB2 clusters were enriched in GO terms for “mitotic spindle assembly checkpoint signaling” and “L-alpha-amino acid transmembrane transport” (P-value < 10^-3^ and FDR < 0.5) (Tables S3m and S3o). Notably, recent studies have suggested that the perturbed cerebral carbohydrate metabolism contributes to AD pathogenesis [65, 66], whereas neuron migration ability contributes to neurogenesis in the brain [67, 68]. These results suggested that the three AD subgroups may differ in these functional features.

We also evaluated the chromosomal distribution of genes belonging to the CA1 or CB1 cluster in N-A1 set, and that of those belonging to the CA2 or CB2 clusters in N-A2 set (Fig. 4). Compared with the chromosomal distribution of all human coding and noncoding genes, both gene sets belonging to the CA1 or CB1 clusters and CA2 or CB2 clusters seemed to be accumulated at Chr. 1, 7, 9, 16, 19, and 22 (Fig. 4).

**Fig. 4.**
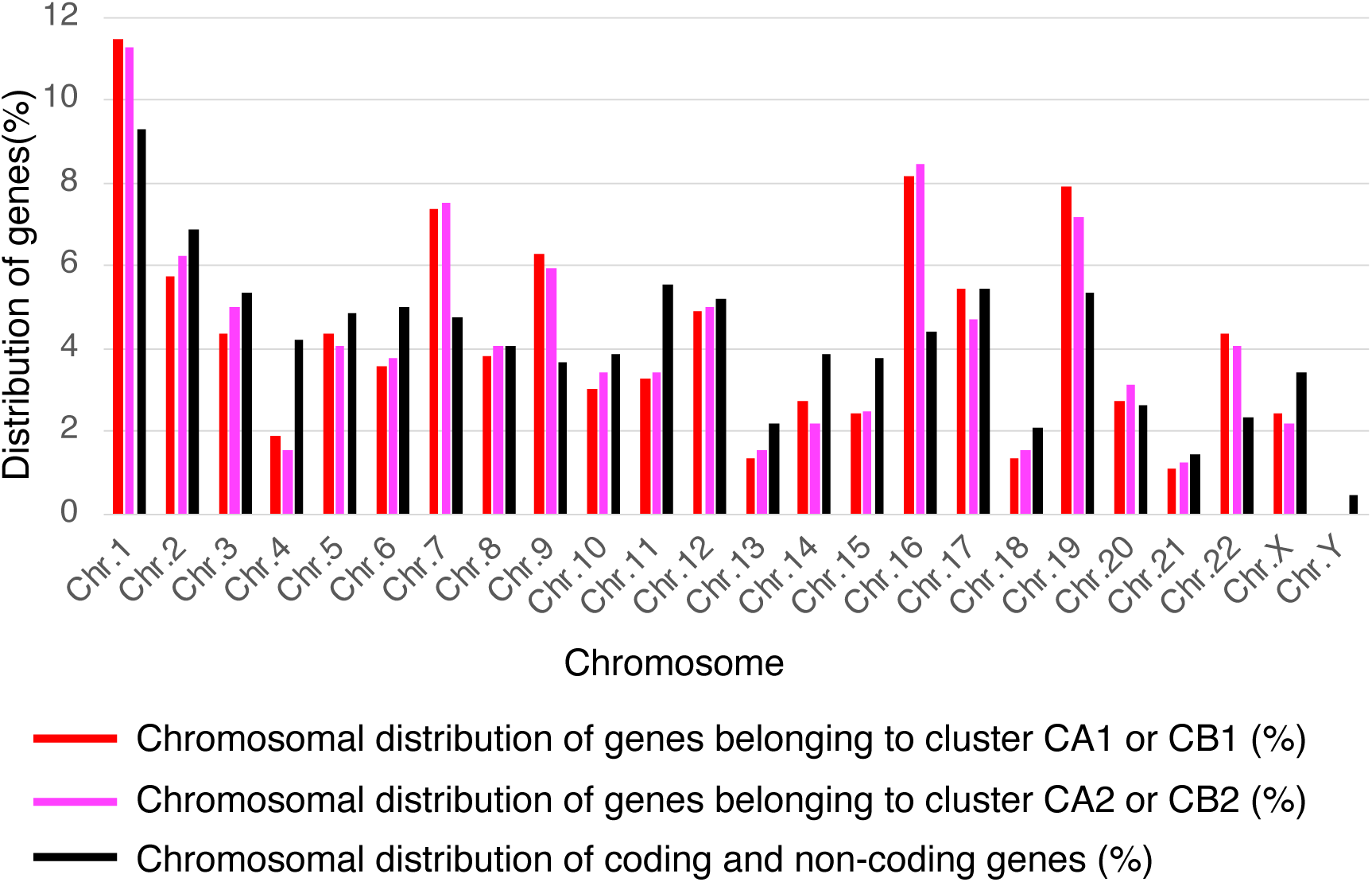
Chromosomal distribution of genes belonging to the CA1 or CB1 cluster (Fig. 3a), and that of genes belonging to the CA2 or CB2 cluster (Fig. 3b). The chromosomal distribution of coding and non-coding genes was estimated using the human genome data GRCh38.

### Epigenome data analysis supported the classification of patients with AD into three groups

The data from the dorsolateral prefrontal cortex of the Rush AD database analyzed in the present study also included information on the genome-wide distribution of epigenomic markers such as histone modifications (H3K4me3, H3K27me3, and H3K27ac) and CTCF binding site distribution for approximately half of all individuals (Table S1). Hence, to classify these individuals based on features other than transcriptome, we next evaluated the similarity of such epigenomic features among individuals using Jaccard coefficients of the genome-wide distribution of previously defined peak regions of epigenome status. At the first divergence of the dendrogram according to hierarchical clustering using Jaccard coefficients, individuals were divided into two groups: a group with a typical epigenome status distribution and another one with an untypical distribution for the respective epigenomic markers (Figs. 5–6 and S4). The Jaccard coefficients of peak regions of epigenome status among individuals suggested that more AD_HL_ individuals exhibited untypical patterns compared with those in other patients with AD (Figs. 5–6 and S4, Table 2); in particular, the AD_HL_ group had a significantly larger number of patients with H3K4me3 modifications compared with the AD_HH_ and AD_HL_ groups (Table 2a). Similar features were obtained for H3K27ac, H3K27me3, and CTCF (Table 2b–2d); however, these results were not significant. Moreover, AD_HL_ individuals tended to exhibit low epigenetic peak region density (Figs. 5–6 and S4).

**Fig. 5.**
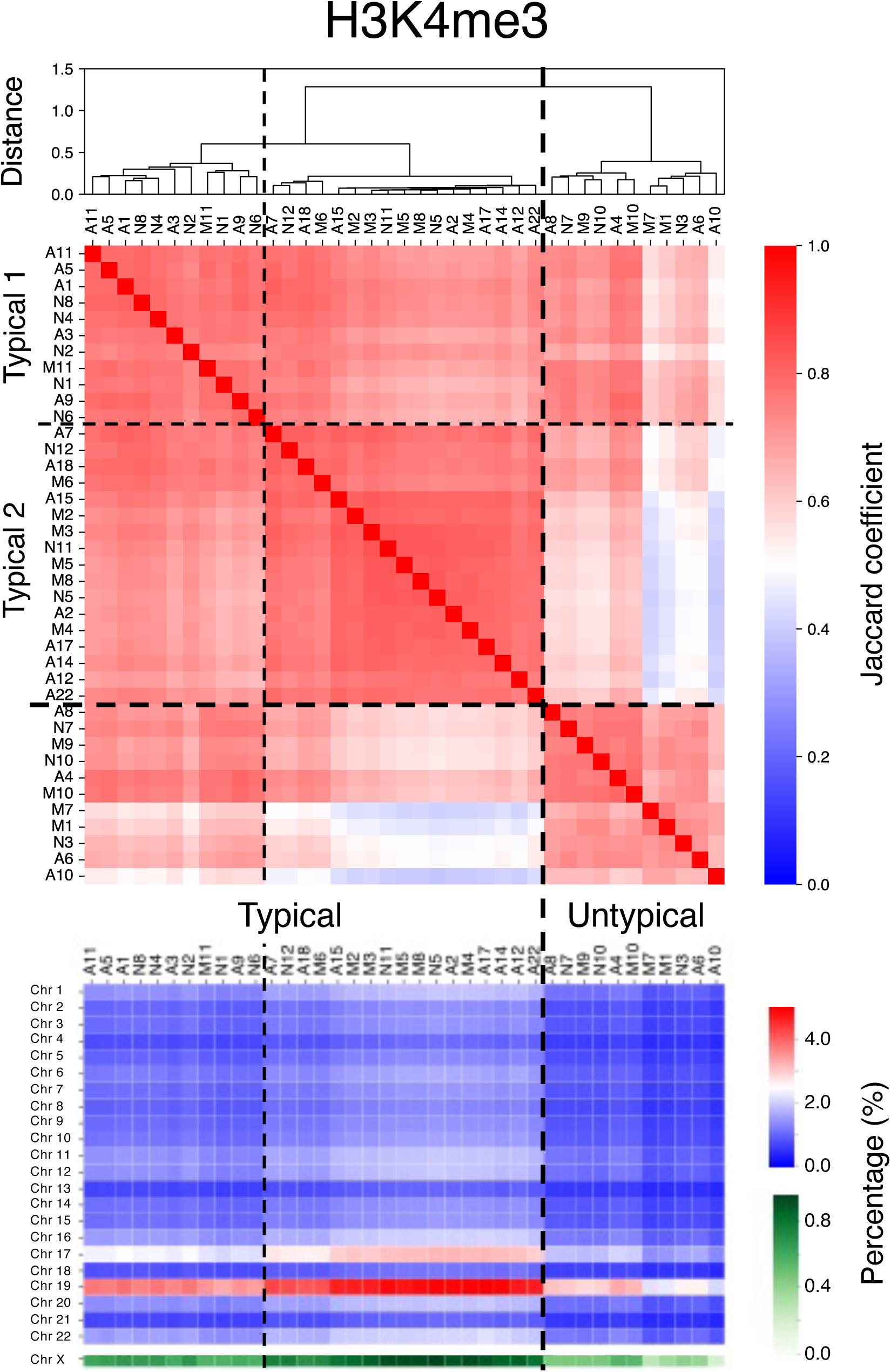
Jaccard coefficient of the genome-wide distribution of H3K4me3 among individuals. Individuals were divided into two groups at the first divergence of the dendrogram. The epigenetic state exhibited by individuals belonging to the group containing the larger number of individuals was regarded as the “typical” epigenetic state, whereas that exhibited by individuals belonging to the group containing the fewer individuals was regarded as the “untypical” epigenetic state. Further, individuals belonging to the “typical” group were divided into two groups, named Typical 1 and Typical 2, at the second divergence of the dendrogram. Dashed lines indicate the boundaries between groups.

**Fig. 6.**
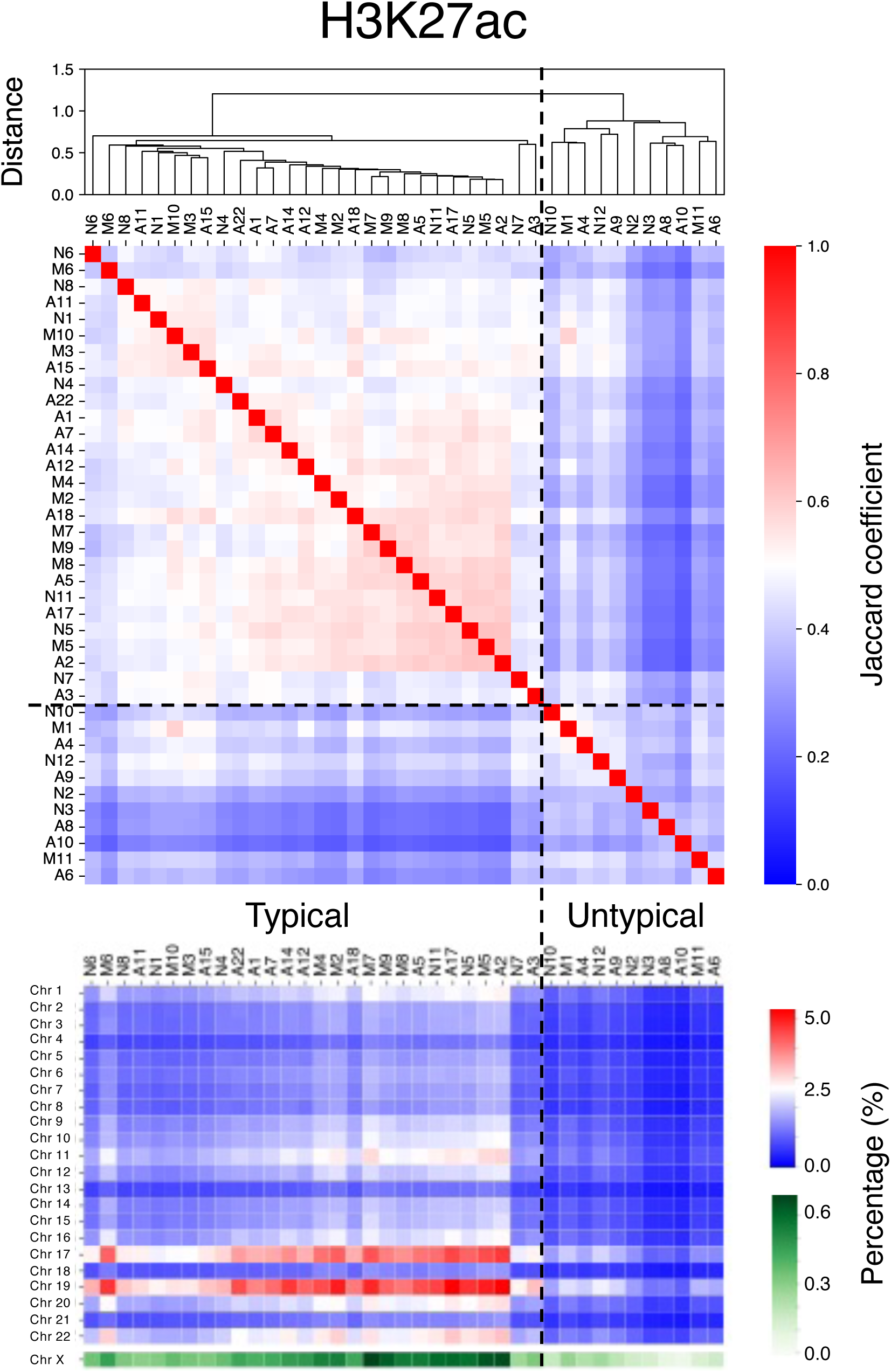
Jaccard coefficient of the genome-wide distribution of H3K27ac among individuals. Analysis was conducted similar to that described in Fig. 5.

**Table 2.** Number of individuals exhibiting the typical genome-wide distributions of each epigenetic state, E_T_, and of those exhibiting untypical distributions, E_U_, among the three AD substages (AD_HH_, AD_LL_, and AD_HL_); H3K4me3 (a), H3K27ac (b), H3K27me3 (c), and CTCF (d). The E_T_ and E_U_ in each substage were obtained from Figures 5, 6, and S4. P-values obtained using the Fisher’s exact test were 0.023843 (a), 0.193439 (b), 0.193439 (c), and 0.285294 (d). Number of individuals exhibiting E_T_ were divided into E_T1_ (Typical 1) and E_T2_ (Typical 2) among the three AD substages by further clustering for H3K4me3 obtained from Figure 5 (e). The P-value obtained using the Fisher’s exact test was 0.0420316 (e).

| H3K4me3 | $AD_{HH}$ | $AD_{LL}$ | $AD_{HL}$ |
| --- | --- | --- | --- |
| # of individuals with typical ($E_T$) | 6 | 4 | 3 |
| # of individuals with untypical ($E_U$) | 0 | 0 | 4 |

(b)
| H3K27ac | $AD_{HH}$ | $AD_{LL}$ | $AD_{HL}$ |
| --- | --- | --- | --- |
| # of individuals with typical ($E_T$) | 5 | 4 | 3 |
| # of individuals with untypical ($E_U$) | 1 | 0 | 4 |

(c)
| H3K27me3 | $AD_{HH}$ | $AD_{LL}$ | $AD_{HL}$ |
| --- | --- | --- | --- |
| # of individuals with typical ($E_T$) | 5 | 4 | 3 |
| # of individuals with untypical ($E_U$) | 1 | 0 | 4 |

(d)
| CTCF | $AD_{HH}$ | $AD_{LL}$ | $AD_{HL}$ |
| --- | --- | --- | --- |
| # of individuals with typical ($E_T$) | 6 | 3 | 4 |
| # of individuals with untypical ($E_U$) | 0 | 1 | 3 |

(e)
| H3K4me3 | $AD_{HH}$ | $AD_{LL}$ | $AD_{HL}$ |
| --- | --- | --- | --- |
| # of individuals with Typical 1 ($E_{T1}$) | 1 | 2 | 2 |
| # of individuals with Typical 2 ($E_{T2}$) | 5 | 2 | 1 |
| # of individuals with untypical (E<sub>U</sub>) | 0 | 0 | 4 |

Further clustering revealed that individuals exhibiting typical H3K4me3 status were further divided into two groups, named Typical 1 and Typical 2 (Fig. 5). Based on their specific epigenome features, more AD_HH_ individuals belonged to the Typical 2 subgroup compared with other patients with AD (Table 3e). These facts suggest that the AD_HH_, AD_HL,_ and AD_LL_ individuals exhibited differences in the genome wide distribution of H3K4me3 modifications.

**Table 3.** Number of APOE2, APOE3, and APOE4 carriers in each clinical diagnostic stage (a), in the two MCI substages (b), and in the three AD substages (c). The genotype of each individual is given in Table S1a and S1b. The P-values obtained using the Fisher’s exact test were 0.8841 (a), 0.08148 (b), and 0.4379 (c).

| Clinical Diagnostic Stage | NCI | MCI | AD |
| --- | --- | --- | --- |
| # of APOE2 carriers | 5 | 3 | 4 |
| # of APOE3 carriers | 16 | 20 | 18 |
| # of APOE4 carriers | 4 | 6 | 6 |

(b)
| MCI substages | MCI<sub>Major</sub> | MCI<sub>NCI_AD</sub> |
| --- | --- | --- |
| # of APOE2 carriers | 1 | 2 |
| # of APOE3 carriers | 18 | 2 |
| # of APOE4 carriers | 5 | 1 |

(c)
| AD substages | AD<sub>HH</sub> | AD<sub>LL</sub> | AD<sub>HL</sub> |
| --- | --- | --- | --- |
| # of APOE2 carriers | 3 | 1 | 0 |
| # of APOE3 carriers | 6 | 7 | 5 |
| # of APOE4 carriers | 1 | 2 | 3 |

### No effect of *APOE* gene polymorphisms in classifying individuals

To examine the relationships between the *APOE* genotype and the subgroup to which individuals belong, the *APOE* genotype of each individual was identified using the RNA-seq read data from the dorsolateral prefrontal cortex of individuals in the Rush AD database. Accordingly, no individuals were found to simultaneously possess the ε2 and ε4 alleles. Therefore, individuals were classified in three categories: APOE2 carrier, possessing one or two ε2 alleles; APOE3 carrier, possessing two ε3 alleles; and APOE4 carrier, possessing one or two ε4 alleles. Notably, the number of APOE2, APOE3, and APOE4 carriers was not correlated with the clinical diagnostic stage of individuals (Tables 3a and S1a–b). In addition, the number of individuals also seemed to be independent of the two MCI (MCI_Major_ and MCI_NCI_AD_) and three AD (AD_HH_, AD_HL_, and AD_LL_) substages (Table 3b–c, Table S1a–b).

### Adjacency network among substages

The data on the expression levels of genes in the M-NA set revealed differences between patients with typical MCI, NCI, and AD, whereas those in the N-A1 or N-A2 sets revealed differences between patients with NCI and AD. To clarify the relationships among disease substages and the pathways of disease progression in these substages, we explored the adjacency networks among substages using M-NA and N-A1 sets, and M-NA and N-A2 sets by performing the partition-based graph abstraction (PAGA) method [69]. Using PAGA, we generated a graph of substages in which the edge weights represented connection confidence. Notably, connection confidence between NCI and AD substages often involved finite values (Table S4); however, the direct connection between NCI and AD substages seemed unnatural in disease progression. Therefore, we assumed that only connections with confidence values sufficiently larger than the maximum confidence values of the connection between NCI and AD substages (MD_N_A_) were effective for the adjacency network among substages (Fig. S5). Subsequently, we used connections with confidence values larger than twice that of MD_N_A_ to construct the adjacency network (Table S4).

Changes in gene expression patterns that characterize disease states are considered to occur gradually as the disease progresses. Therefore, we observed that disease appeared to progress from NCI to AD along the previously obtained adjacency network. Note that the adjacency network obtained using the M-NA and N-A1 sets (Fig S5a) and that constructed using the M-NA and N-A2 sets (Fig. S5b) exhibited a slightly different structure because of differences in the contained genes in these sets. Thus, we focused on the network constructed using common connections obtained from both networks constructed using the M-NA and N-A1 sets (Fig S5a) and M-NA and N-A2 sets (Fig. S5b) (Fig. 7). We did not obtain any connections from NCI to MCI_Major_, the major group of MCI individuals, in any of the networks (Figs. 7 and S5a–b), which seemed unnatural for AD progression. Thus, we performed more detailed subgroup classifications of patients with NCI, MCI, and AD to reveal the relationship between NCI and MCI_Major_ individuals.

**Fig. 7.**
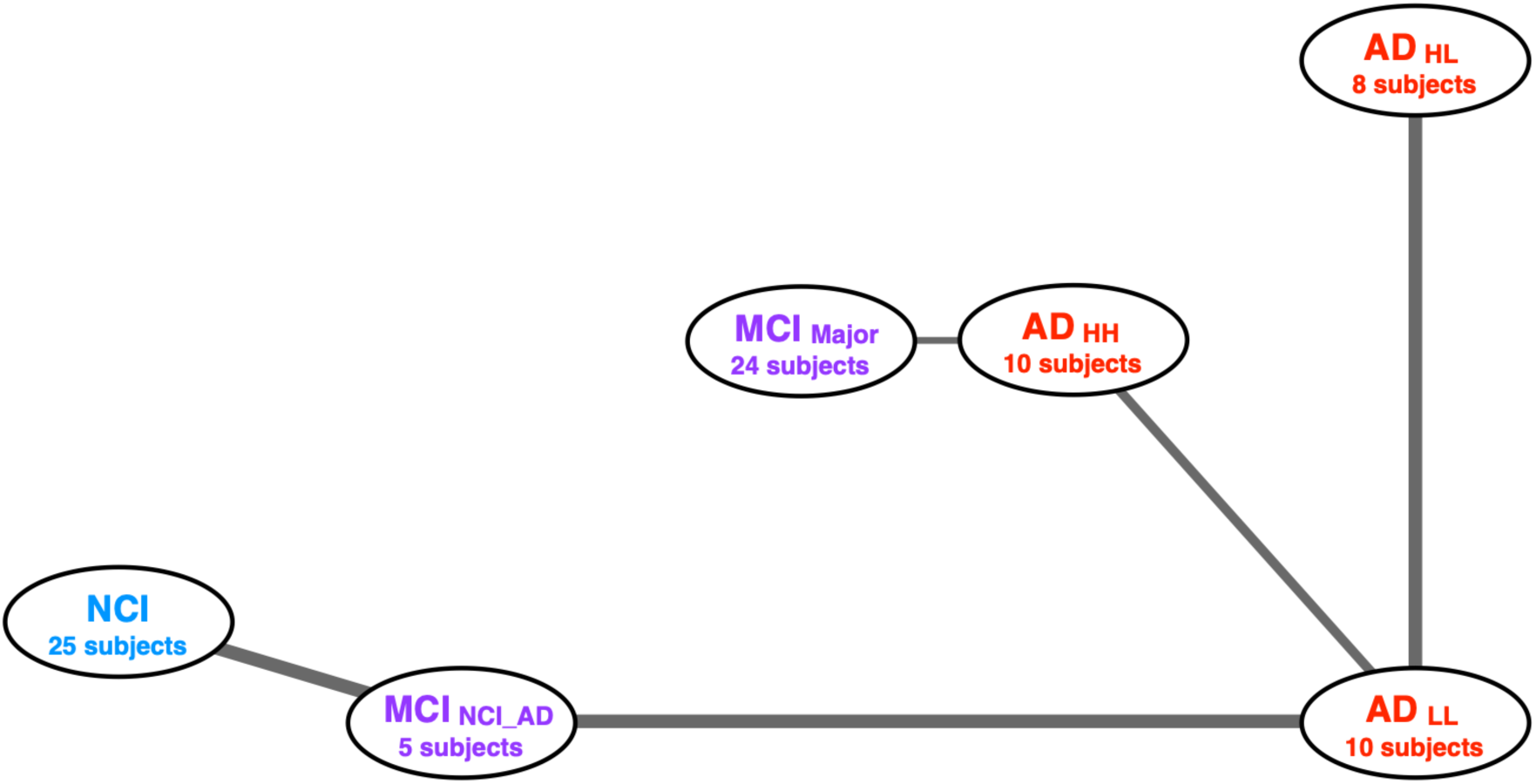
Adjacency network among substages. The network constructed using common connections obtained from both the network constructed using the M-NA and N-A1 sets (Fig S5a) and that constructed using the M-NA and N-A2 sets (Fig. S5b). The width of each edge indicates the connection confidence between each pair of connected nodes.

### Detailed classification of patients with MCI by comparing between patients with NCI and those with MCI and between patients with MCI and those with AD

To classify patients with MCI more comprehensively, we conducted comparisons between patients with NCI and those with MCI and between patients with MCI and those with AD. First, we performed hierarchical clustering of individuals using the genes satisfying the following criteria: the mean expression level (TPM) was higher in the MCI group than that in the NCI group with smaller P-values in the likelihood ratio test than the P^D^_MN_ threshold or the mean expression level (TPM) was lower in the MCI group than that in the NCI group, with smaller P-values in the likelihood ratio test than the P^D^_NM_ threshold. We performed cluster analysis using each gene set within the range P^D^_NM_ ≤ 0.05 and P^D^_MN_ ≤ 0.05. However, we did not identify any cases in which patients with NCI were completely separated from those with MCI (Figs. 8a and S6a). However, clusters with significantly more patients with NCI (NCI clusters) and MCI clusters were formed within all cases of P^D^_NM_ and P^D^_MN_ except in the case of P^D^_NM_ = 0.05 and P^D^_MN_ = 0.01 (Fig. 8a).

**Fig. 8.**
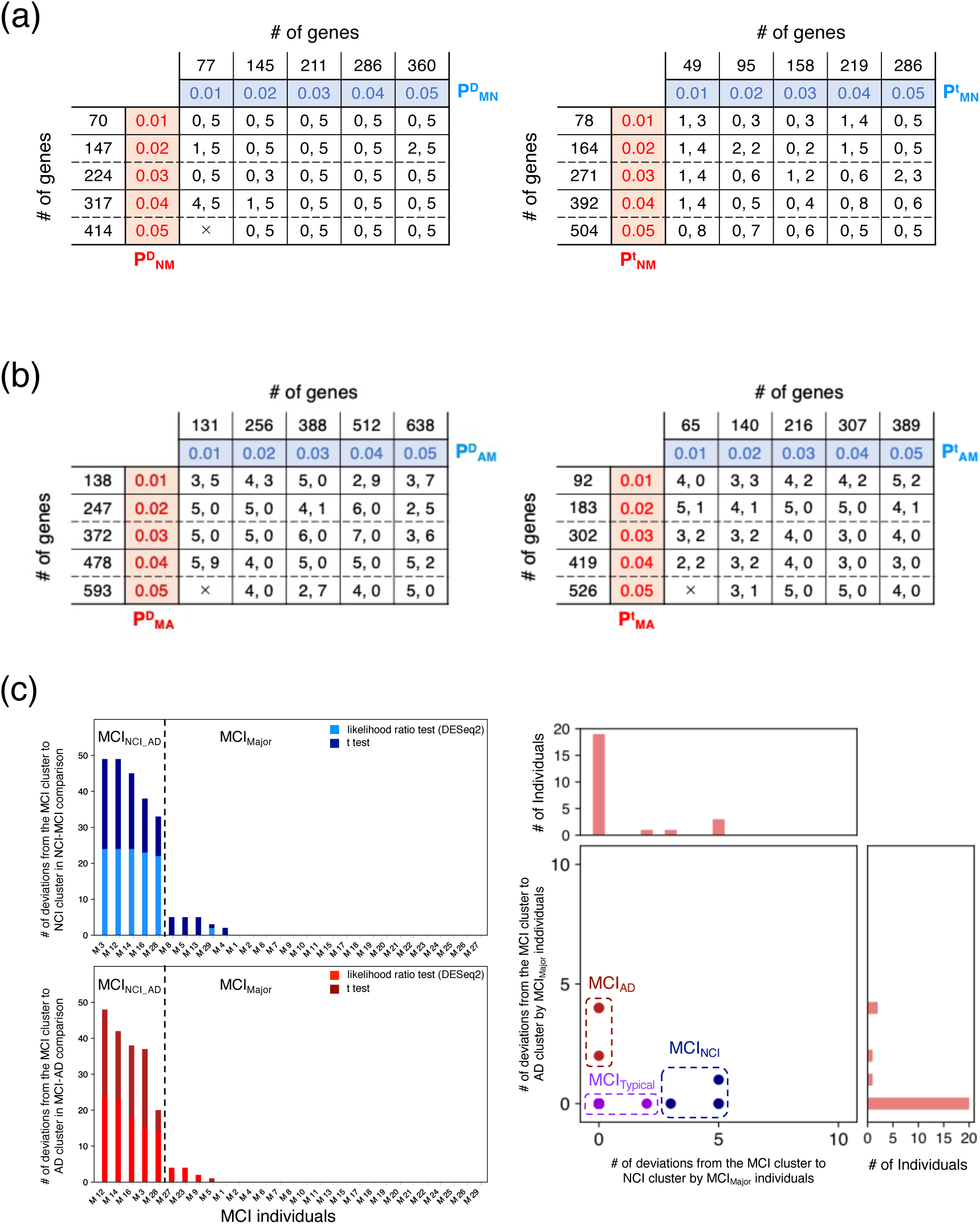
Detailed classification of patients with MCI. Number of patients with NCI who belong to the MCI cluster (left value) and that of patients with MCI who belong to the NCI cluster (right value) as a function of P^D^_NM_ and P^D^_MN_ (a, left), and as a function of P^t^_NM_ and P^t^_MN_ (a, right). Number of patients with MCI who belong to the AD cluster (left value) and that of patients with AD who belong to the MCI cluster (right value) as a function of P^D^_MA_ and P^D^_AM_ (b, left), and as a function of P^t^_MA_ and P^t^_AM_ (b, right). Number of deviations of each patient with MCI from the MCI to the NCI (c, upper left) and AD (c, lower left) cluster. Broken lines indicate boundaries between MCI_NCI_AD_ and MCI_Major_ individuals. Scatter plot of MCI_Major_ individuals as a function of the number of times that each individual belonged to the NCI and AD cluster (c, right). Using k-means clustering (Fig. S6c), MCI_Major_ individuals were divided into MCI_Typical_, MCI_NCI_, and MCI_AD_ groups.

When we used two-tailed *t*-tests instead of the likelihood ratio test to determine the P-values, our results were qualitatively the same. P^t^_MN_ and P^t^_NM_ were defined instead of P^D^_MN_ and P^D^_NM_, respectively. Analysis using each gene set within the ranges P^t^_MN_ ≤ 0.05 and P^t^_NM_ ≤ 0.05 revealed no cases in which patients with NCI were completely separated from those with MCI (Figs. 8a and S6a). However, NCI and MCI clusters were formed within all cases of P^t^_NM_ and P^t^_MN_ (Fig. 8a).

Next, we performed the same analyses for patients with MCI and those with AD. In this case, P^D^_MA_, P^D^_AM_, P^t^_MA_, and P^t^_AM_ were given as values corresponding to P^D^_NM_, P^D^_MN_, P^t^_NM_, and P^t^_MN_, respectively. Similar to the above results, we did not identify any cases in which patients with MCI were completely separated from those with AD (Fig. 8b and S6b). However, MCI and AD clusters were formed within all cases of P^D^_MA_, P^D^_AM_, P^t^_MA_, and P^t^_AM_ except in the case of P^D^_MA_ = 0.05 and P^D^_AM_ = 0.01 and that of P^t^_MA_ = 0.05 and P^t^_AM_ = 0.01 (Fig. 8b).

In this study, variations were observed in patients with MCI who deviated from the MCI cluster depending on P^D^_NM_, P^D^_MN_, P^t^_NM_, P^t^_MN_, P^D^_MA_, P^D^_AM_, P^t^_MA_, and P^t^_AM_ values. For each patient with MCI, we counted the number of times they belonged to the NCI cluster upon changing the P^D^_NM_, P^D^_MN_, P^t^_NM_, and P^t^_MN_ values, and those belonging to the AD cluster upon changing the P^D^_MA_, P^D^_AM_, P^t^_MA_, and P^t^_AM_ values (Fig. 8c). Consequently, individuals M3, M4, M5, M8, M12, M13, M14, M16, M28, and M29 belonged to the NCI cluster more than once, whereas individuals M3, M5, M9, M12, M14, M16, M23, M27, and M28 belonged to the AD cluster more than once.

Notably, individuals M3, M12, M14, M16, and M28, (same as those in MCI_NCI_AD_), clearly exhibited a greater number of deviations from patients with NCI and AD than MCI_Major_ individuals (Fig. 8c). To classify MCI_Major_ individuals in more detail, we performed k-means clustering using the number of times each individual belonged to the NCI cluster and that belonging to the AD cluster (Figs. 8c and S6c). As a result, MCI_Major_ individuals were divided into three subtypes: MCI_Typical_ individuals (17 individuals) who rarely or never belonged to other clusters; MCI_NCI_ individuals (4 individuals) who may belong to NCI clusters only; and MCI_AD_ individuals (3 individuals) who may belong to AD clusters only (Table S1a–S1b). Thus, MCI was divided into four substages: MCI_Typical_, MCI_NCI_, MCI_AD_, and MCI_NCI_AD_.

### Detailed classification of patients with NCI by comparing them with patients with MCI

Similar analysis showed that individuals N11, N12, N18, N19, N21, and N22 belonged to the MCI cluster more than once when the P^D^_NM_, P^D^_MN_, P^t^_NM_, and P^t^_MN_ values were changed (Fig. 9a). After k-means clustering of patients with NCI using the number of times each individual belonged to the MCI cluster, patients with NCI were divided into two clusters: one comprising N11, N21, and N22, and another comprising those who never belonged to the MCI cluster (Figs. 9a and S7a). Therefore, these individuals were classified as typical patients with NCI belonging to the NCI cluster, named NCI_Typical_ individuals, who were assumed to be in the NCI_Typical_ substage. Individuals N12, N18, and N19 were classified as patients with NCI close to MCI, named NCI_MCI_ individuals, who were assumed to be in the NCI_MCI_ substage (Table S1a–S1b).

**Fig. 9.**
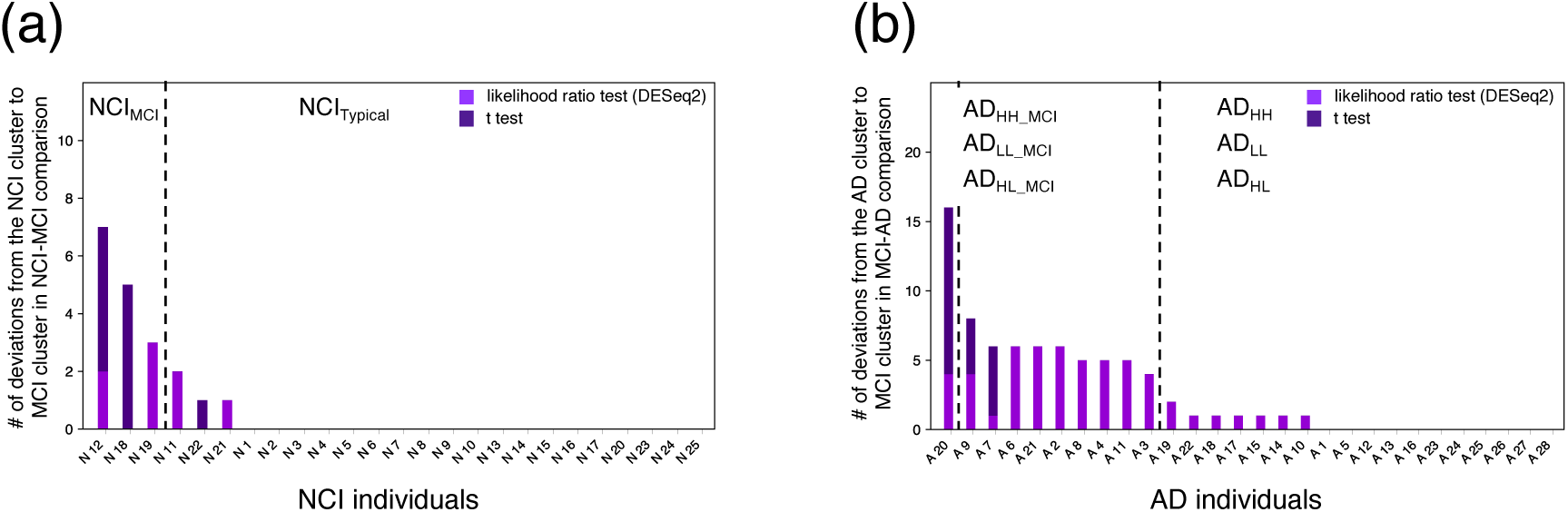
Detailed classification of patients with NCI. Number of deviations of each patient with NCI from the NCI to the MCI cluster (a), and that of each patient with AD from the AD to the MCI cluster (b). Broken lines indicate boundaries among clusters of individuals obtained via k-means clustering (Fig. S7a and S7b).

### Detailed classification of patients with AD by comparing them with patients with MCI

Similarly, 17 patients with AD belonged to the MCI cluster more than once, when the P^D^_MA_, P^D^_AM_, P^t^_MA_, and P^t^_AM_ values were changed (Fig. 9b). Using k-means clustering of patients with AD based on the number of times each individual belonged to the MCI cluster, patients with AD were divided into three clusters, with individuals A10, A14, A15, A17, A18, A19, A22, and other patients with AD who never belonged to the MCI cluster being included in the same cluster (Figs. 9b and S7b). Therefore, these individuals were expected to be in one of the typical AD substages: AD_HH_, AD_HL_, or AD_LL_ (Table S1a–S1b). In contrast, individuals A2, A3, A4, A6, A7, A8, A9, A11, A20, and A21 were assumed to be at one of the AD substages close to MCI and were named AD_HH_MCI_, AD_HL_MCI_, or AD_LL_MCI_ (Table S1a–S1b).

### Adjacency network among substages obtained after detailed classifications of individuals suggests multiple disease progression pathways from NCI to AD

Detailed classifications of individuals suggested that patients with NCI were classified into NCI_Typical_ and NCI_MCI,_ patients with MCI were classified into MCI_Typical_, MCI_NCI_, MCI_AD_, and MCI_NCI_AD,_ and patients with AD were classified into AD_HH_, AD_HH_MCI_, AD_HL_, AD_HL_MCI_, AD_LL_, and AD_LL_MCI_. We constructed adjacency networks among these 12 substages using gene expression data from the M-NA and N-A1 sets (Fig. S8a, Table S5a) as well as from the M-NA and N-A2 sets (Fig. S8b, Table S5b) as previously described. Subsequently, we focused on the adjacency network that consisted of common connections between the network from the M-NA and N-A1 sets, and that from the M-NA and N-A2 sets (Fig. 10). We identified more than four progression paths from NCI to AD that passed through different substages, as follows: I) from NCI (NCI_Typical_ or NCI_MCI_) to AD_LL_ through MCI_NCI_AD_, II) from NCI to MCI_NCI_ and from MCI_NCI_ to AD_LL_ through AD_HH_MCI_ and AD_LL_MCI_, respectively, III) from NCI to MCI_NCI_ and from MCI_NCI_ to AD_HH_ through MCI_Typical_ or AD_HH_MCI_, and IV) from NCI to AD_HL_ or AD_LL_ via MCI_NCI_, MCI_AD_, or AD_HL_MCI_. This suggested that at least four disease progression pathways are active in the three classified AD substages.

**Fig. 10.**
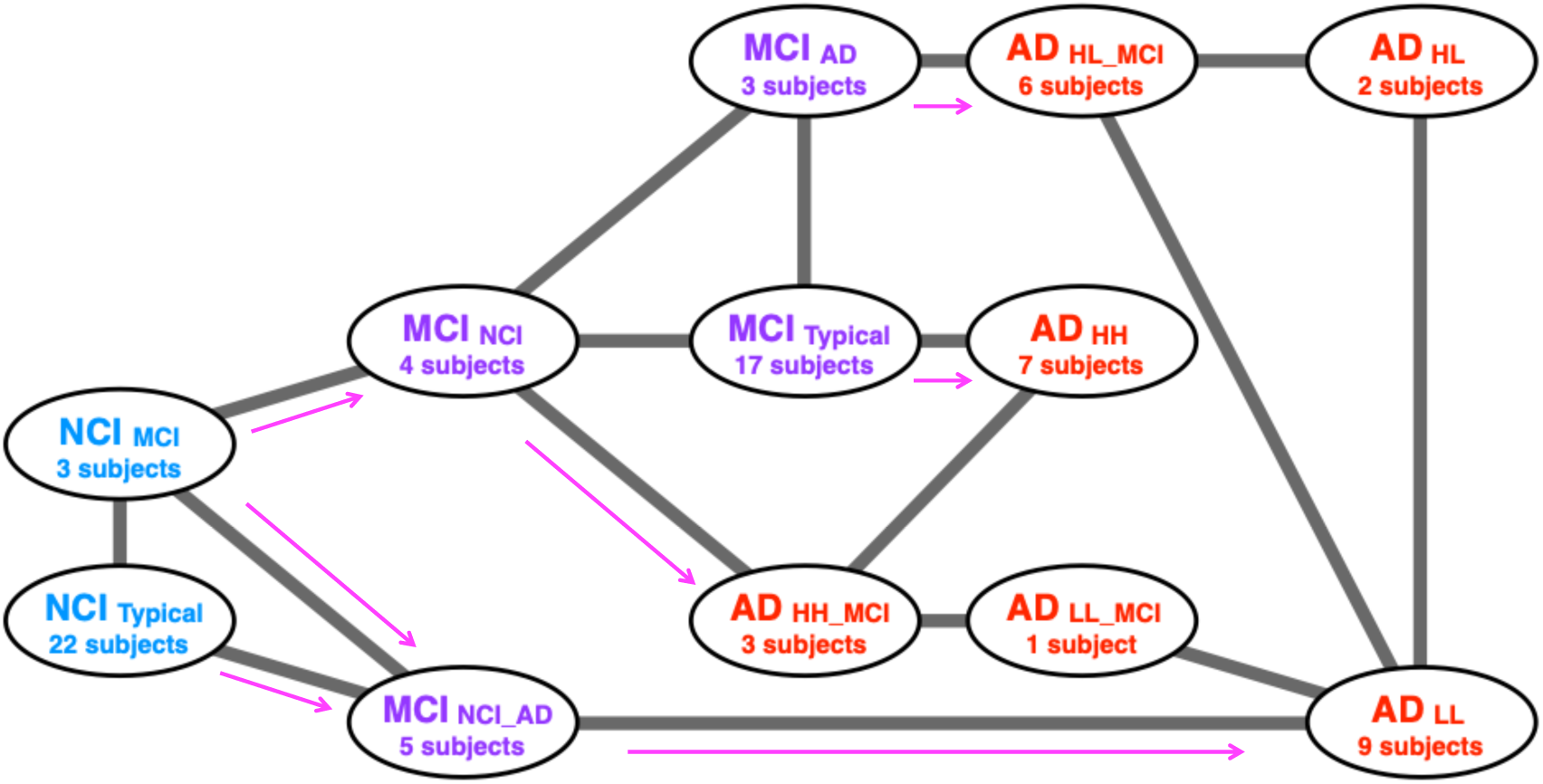
Adjacency networks among substages. A network containing common connections from both networks obtained using the data in the M-NA and N-A1 sets (Fig. S8a) and that in the M-NA and N-A2 sets (Fig. S8b). The width of each edge indicates the connection confidence between each pair of connected nodes. Arrows indicated the inferred direction of disease progression.

### Comparisons with the findings of recent studies

Many recent studies have performed transcriptome analyses and identified various significant differentially expressed genes (significant DEGs) between healthy (control) individuals and patients with AD, which they termed “AD risk genes” [50, 59-61]. The cluster analysis of patients with NCI, MCI, and AD and that of patients with NCI and AD was performed using these AD risk genes (Fig. S9, Table S6). However, these results differed from the individual clustering results in Figures 1, 2, and 3. Notably, most of the AD risk gene data used here were obtained from research on the hippocampus, whereas the data analyzed in the previous sections of this study were obtained from the prefrontal cortex. A recent study showed that gene expression patterns differ between the hippocampus and prefrontal cortex [39]. Hence, the observed inconsistencies were attributed to these differences.

From the dataset analyzed in the previous sections, no significant DEGs were detected between patients with NCI and those with AD, whereas only nine significant DEGs were identified among patients with NCI, MCI, and AD (Tables S1a–S1b and S7). Herein, significant DEGs were evaluated via the likelihood test using DEseq2 with FDR < 0.01. Notably, the abovementioned study reported that not many significant DEGs were yielded in areas belonging to the prefrontal cortex [39]. These facts supported the consistency between the results of the present and those of recent studies.

## DISCUSSION

In this study, the analysis of data from women individuals in the Rush AD database suggested that the pathological progression pathway of AD is nonuniform, exhibiting different typical forms of deterioration. First, cluster analysis using gene expression data from individuals diagnosed with NCI, MCI, and AD suggested that NCI can be classified in at least two substages, whereas MCI and AD can be classified in four and six substages, respectively. Differences among AD substages were identified based on epigenomic features, in particular the genome-wide distribution of H3K4me3. However, the *APOE* gene polymorphisms of individuals appeared to not correlate with the disease substage. In addition, some of the genes and gene sets that were suggested in recent studies to be AD-associated contributed to the characterization of each AD substage in the present study. Second, an inference of adjacency networks among the different substages suggested the existence of four typical NCI → AD disease progression pathways from different MCI to different AD substages. These findings on disease progression features will facilitate further advancements in AD research. Moreover, they are expected to induce changes in the strategic bases of AD prevention, treatment, and studies for biomarkers [70-72].

Recent studies using hippocampal and frontal lobe transcriptomes have also suggested the existence of multiple AD subtypes. However, the relationship between multiple states, such as their connection in a series on the same disease progression path or on different paths, has not been clarified. Furthermore, these studies did not focus on the diversity of NCI and MCI. In contrast, in this study, both patients with NCI and MCI were divided into multiple subgroups at different substages based on their respective gene expression characteristics. In addition, estimating the connectivity among substages clearly revealed the existence of multiple pathological pathways of disease progression.

This study analyzed data from the dorsolateral prefrontal cortex obtained from the Rush AD database, rather than the hippocampus, which has been the focus of many previous studies. The reason for analyzing this dataset in this study was that, it included not only transcriptome data but also data on the genome-wide distribution of histone modifications (H3K4me3, H3K27me3, H3K27ac) and CTCF binding sites (Table S1). Such datasets can potentially reveal the relationship between a wide range of transcriptional regulatory states and disease conditions. Recently, by analyzing such datasets, changes in CTCF binding and transcription activities were shown to be associated with changes in pathological conditions [73]. The analysis of such epigenome data in the present study suggests that the genome-wide distribution of H3K4me3 differed among individuals belonging in three different AD substages. However, the mechanism by which such differences in the epigenetic status are related to gene expression patterns remains unclear and warrants elucidation in future studies.

Notably, many recent studies have performed transcriptome analyses to identify significant DEGs between healthy individuals and patients with AD. However, by focusing on the expression patterns of more widespread candidate genes rather than the significant DEGs, we identified different characteristics appearing in the dorsolateral prefrontal cortex between healthy individuals and patients with AD and that disease substages occur in response to changes in the disease state.

In addition, many studies have focused on gene expression patterns in the hippocampus. One such previous research suggested the existence of multiple states in the hippocampus of patients with AD based on differences in the gene expression profiles [39]. However, as previously mentioned, the gene expression patterns differ between the hippocampus and prefrontal cortex, and these differences exhibit further variations between healthy individuals and patients with AD. Hence, different results would be obtained even from the same analysis. Importantly, the hippocampal data analyzed in previous studies can be analyzed in the same way as in this study; connections between substages can be extracted and pathological routes can be estimated, all of which are crucial issues that need to be addressed in the future.

The present study has some limitations. We only classified female individuals as we used transcriptome and epigenomic data of female patients from the Rush AD database. However, the insights of this study are expected to be useful not only for female patients but also for understanding the rapid progression of symptoms in male patients. To validate this expectation, the dataset from male individuals should also be analyzed. In this dataset, however, the number of female patients in the NCI, MCI, and AD groups was 25, 29, and 28, respectively, whereas that of men was 14, 4, and 7, respectively, that is, three times smaller than that of female individuals (seven and four times smaller than those of female patients with MCI and AD). Moreover, the number of male patients with MCI and AD in the database (four and seven) was smaller than that of individuals in each of the three AD groups obtained for female individuals. With such restricted data, performing similar analyses on male individuals or sex-independent analyses was impossible. Obtaining more data on male patients is one of the essential issues that must be resolved in the future for performing such analyses and eliminating potential sex-related bias.

The likelihood ratio test using DESeq2 and *t*-test were used to evaluate the differences in RNA-seq data between different groups when searching for gene sets to characterize and classify individuals. The *t*-test is generally better for detecting differences and was expected to identify more candidate genes in a gene set than the likelihood ratio test or other nonparametric testing methods. However, the *t*-test produces more false positives than other testing methods when data deviate from a Gaussian distribution. The gene sets used in this study were expected to contain few false-positive genes because they were chosen under imposing the additional condition that healthy individuals could be clearly distinguished from patients through cluster analysis. However, inappropriate genes may have still been retained. For more precise consideration, verification is necessary using various nonparametric tests.

In addition to the transcriptome of each individual, the database targeted for the present analysis also contained various epigenome data; however, other information on each individual, such as postmortem interval, brain PH, and medical history was not included. Therefore, although this study has yielded some unprecedented findings, the validation and characterization of each substage through comparisons with other physiological data could not be included. However, such analysis may be achieved by combining meta-analysis with databases focusing on other feature groups.

## EXPERIMENTAL PROCEDURES

### Data processing

Gene expression data (read count and TPM) from the dorsolateral prefrontal cortex of patients with NCI, MCI, and AD were acquired from the Rush AD database. In the present analysis, the data of genes with average expression levels (TPM) in patients with NCI, MCI, and AD were smaller than 0.1, whereas those of spike-in-RNAs were eliminated.

The data provided by the Rush AD study in ENCODE does not publish information on RNA quality such as the postmortem interval of samples and RIN values. However, according to ENCODE standards, no specific alerts regarding the quality of the RNA-seq data used in this study were provided. Thus, the quality of the data was assumed to be guaranteed. The set of genes whose expression changes between patients with NCI and AD and among patients with NCI, MCI, and AD, involved many genes identified as AD risk genes in previous studies. This further supported the lack of data bias that would contradict previous studies.

### Statistical analyses

Two-tailed *t*-tests and ANOVA were performed using Python (version 3.9.16). All *t*-tests in this study were Welch’s *t*-tests. The false discovery rate (FDR) was estimated through a set of P-values using the Benjamini–Hochberg procedure [74] implemented in MultiPy (https://puolival.github.io/multipy/) [75]. The search for DEGs via the likelihood ratio test using DESeq2 and the calculation of P-values for the Fisher’s exact test were performed using R (version 4.1.3).

### Cluster analyses

Cluster analyses were performed using the values of X = log_2_ (TPM + 1) – C, where C was chosen to satisfy the average of X over individuals = 0 for each gene. iDEP.96 (http://bioinformatics.sdstate.edu/idep96/) was used for hierarchical clustering [76]. In hierarchical clustering, the group-average method, was used to create a dendrogram, using the correlation coefficient as the distance. If the cluster formed by the first branch of hierarchical clustering satisfied the following criteria for a certain group (NCI, MCI, and AD), then that cluster was regarded as a cluster including significantly more individuals belonging to that group. These criteria were as follows: i) containing more than 2/3 of the individuals belonging to that group, and ii) Fisher exact test resulting in a P-value < 0.05. Further, k-means cluster analysis was performed using the Python package scikit-learn (version 1.0.2) [77]. The number of clusters in each k-means cluster analysis was determined using the elbow method.

The construction of UMAP was performed using sc.tl.pca() [77] with default parameters, scanpy.pp.neighbors() with “n_neighbors” = 30 [78], and the scanpy.tl.umap() functions with n_components = 3 [69] of scanpy (version 1.9.3) according to the literature on basic workflows for clustering analysis using Scanpy (https://scanpy-tutorials.readthedocs.io/en/latest/pbmc3k.html). All principal components were used for clustering using the scanpy.pp.neighbors() function. The obtained results were qualitatively independent of the parameters of these applications.

### Epigenomic data analysis

The Jaccard coefficient of the genome-wide distribution of epigenomic markers among individuals was estimated as follows. The bigwig formatted files of fold change over control of these epigenome markers were acquired from individuals whose ChIP-seq data of H3K4me3, H3K27ac, H3K27me3, and CTCF were provided in the Rush AD database. From each bigwig format data, bedgraph format data was obtained using bigWigToBedGraph [79]. From the bedgraph format data, bed format data of peak regions were obtained following the removal of regions with FC < 5. The Jaccard coefficients between each pair of individuals were estimated using Bedtools (version 2.31.0). The density of each epigenomic marker on each chromosome was estimated according to the following equation: [Total length of genome regions with narrow peaks in ChIP-seq data in the chromosome obtained from the bed formatted file (bp)]/[Total length of chromosome obtained from GRCh38 (bp)].

### Analysis of *APOE* gene polymorphisms

RNA-seq read data (fastq format data) from the dorsolateral prefrontal cortex of patients with NCI, MCI, and AD were acquired from the Rush AD database. Fastp (https://github.com/OpenGene/fastp) was used to remove low-quality reads and trim adapter sequences. Following fastp processing, the reads were mapped to the CDS sequence of the *APOE* exon 4 of GRCh38 using Hisat2 (https://daehwankimlab.github.io/hisat2/), and the results were obtained as a bam format file. Variant calling was performed using the obtained bam format file and CDS sequence data of the GRCh38 *APOE* exon 4 using bcftools (https://github.com/samtools/bcftools). After filtering out low-quality calls (QUAL<20), genotypes were determined based on the type of mutation and allele frequency.

### Enrichment analysis

Enrichment analysis was performed for the M-NA, N-A1, and N-A2 sets using Metascape (v3.5.20240101) (https://metascape.org/gp/index.html#/main/step1) [80]. The biological process terms in Gene Ontology were the main focus of this study.

### Construction of an adjacency network among substages

The adjacency network among substages was estimated via PAGA [69] using the gene expression data of each individual. Each group at each substage was regarded as a partition in PAGA. PAGA can generate a graph of substages in which the edge weights represent connection confidence. PAGA was performed using sc.tl.pca() [77] with default parameters, scanpy.pp.neighbors() with “n_neighbors” = 2 [78], and the scanpy.tl.paga() functions [69] of scanpy (version 1.9.3) according to the literature for trajectory inference using transcriptome data (https://scanpy-tutorials.readthedocs.io/en/latest/paga-paul15.html). All principal components were used for clustering using the scanpy.pp.neighbors() function. Additionally, to extract the essential network among substages, the value of the parameter “n_neighbors” of the function scanpy.pp.neighbors() was set as two, which is the minimum value in the range of this parameter. Thus, a network with nonredundant connections among substages was expected to be obtained.

## AUTHOR CONTRIBUTIONS

Kousei Honda and Akinori Awazu conceived and designed the study. Kousei Honda performed the analyses. Kousei Honda and Akinori Awazu drafted the manuscript.

## ACKNOWLEDGEMENTS

This research was supported by JST Moonshot R&D (Grant Number JPMJMS2021) and a JSPS KAKENHI grant (award number 21K06124, 24K09248). Computations were partially performed on the NIG supercomputer at the ROIS National Institute of Genetics.

## Notes

### Competing Interest Statement

The authors have declared no competing interest.

### Summary of Updates

The title was revised. Some results were added.

