## Supplementary material for "Potential multiple disease progression pathways in female patients with Alzheimer’s disease inferred from transcriptome and epigenome data of the dorsolateral prefrontal cortex": H_A_sup.docx

**SUPPORTING INFORMATION**

**Table S1.** Detailed features of each individual (a-b). The name, ID in the ENCODE database, age, diagnostic stage, inferred substage, inferred detailed substage, first, second, and third quantiles of MOR, availability of epigenome data, *APOE* genotype estimated using RNA-seq data, and gene expression levels (TPM (a), and read count (b)) of each individual are included. Age 90+ indicates that the individual is aged > 90 years. The results of the two-tailed *t*-test and DEseq2, which were used to examine the difference in gene expression levels between individuals < 85 years and those > 90 years, are also shown. Total number of reads from ChIP-seq data for H3K27ac, H3K27me3, H3K4me3, and CTCF (c) mapped to the human genome (GRCh38).

(a) “Table_S1a.xlsx”

(b) “Table_S1b.xlsx”

(c) “Table_S1c.xlsx”

**Table S2**. Detailed results of the clustering of individuals using the M-NA set (a). The name, age, diagnostic stage, inferred substage, inferred detailed substage, and expression levels (TPM) of genes of each individual are shown. The order of genes was the same as that in Figure 1b. Detailed results of the enrichment analysis were obtained using all genes in the M-NA set (b), those with higher expression levels in MCI_Major_ individuals (c), and AD risk genes with lower expression levels in MCI_Major_ individuals (d).

(a) “Table_S2a.xlsx”

(b) “Table_S2b.xlsx”

(c) “Table_S2c.xlsx”

(d) “Table_S2d.xlsx”

**Table S3**. List of genes in the N-A_D1_ set and their expression levels (a). List of genes in the N-A_D2_ set and their expression levels (b). List of genes in the N-A_t_ set and their expression levels (c). List of genes in the N-A1 set and their expression levels (d). List of genes in the N-A2 set and their expression levels (e). The order of genes in Tables d and e are the same as that in Figure 2c and 2d, respectively. Genes in clusters CA1, CB1, CA2, and CB2 are also indicated. The name, age, diagnostic stage, inferred substage, inferred detailed substage, and expression levels (TPM) are included. The P-values of ANOVA with the result of the Benjamini–Hochberg procedure (FDR < 0.1) of the expression level of each gene when the NIC stage is divided into three or four subgroups, and that of each gene among the three subgroups of AD, are also shown. Detailed results of the enrichment analysis were obtained using the genes in the N-A1 set (f), genes in the N-A1 set with higher expression levels in AD than in NCI (g), genes in the N-A1 set with lower expression levels in AD than in NCI (h), genes in the N-A2 set (i), genes in the N-A2 set with higher expression levels in AD than in NCI (j), and genes in the N-A2 set with lower expression levels in AD than in NCI (k). Detailed results of the enrichment analysis obtained using genes in clusters CA1 (l), CB1 (m), CA2 (n), and CB2 (o).

(a) “Table_S3a.xlsx”

(b) “Table_S3b.xlsx”

(c) “Table_S3c.xlsx”

(d) “Table_S3d.xlsx”

(e) “Table_S3e.xlsx”

(f) “Table_S3f.xlsx”

(g) “Table_S3g.xlsx”

(h) “Table_S3h.xlsx”

(i) “Table_S3i.xlsx”

(j) “Table_S3j.xlsx”

(k) “Table_S3k.xlsx”

(l) “Table_S3l.xlsx”

(m) “Table_S3m.xlsx”

(n) “Table_S3n.xlsx”

(o) “Table_S3o.xlsx”

**Table S4**. Matrix of confidence regarding the connections among patients with NCI, MCI_Major_, MCI_NCI_AD_, AD_HH_, AD_HL_, and AD_LL_ individuals estimated from the expression levels of the M-NA and N-A1 sets (a), and M-NA and N-A2 sets (b).

(a) “Table_S4a.xlsx”

(b) “Table_S4b.xlsx”

**Table S5**. Matrix of confidence regarding the connections among the NCI_Typical_, NCI_MCI_, MCI_Typical,_ MCI_NCI_, MCI_AD_, MCI_NCI_AD_, AD_HH_MCI_, AD_HL_MCI_, AD_LL_MCI_, AD_HH_, AD_HL_, and AD_LL_ individuals estimated from the expression levels of the M-NA and N-A1 sets (a), and M-NA and N-A2 sets (b).

(a) “Table_S5a.xlsx”

(b) “Table_S5b.xlsx”

**Table S6**. Detail results of the cluster analysis of patients with NCI, MCI and AD (a) and that of patients with NCI and AD (b) using AD risk genes previously reported in the literature (Hu et al. 2017; Xiang et al. 2018; Rahman et al. 2019; Yang et al. 2022). The name, age, diagnostic stage, inferred substage, inferred detailed substage, and expression levels (TPM) of genes of each individual are shown. Many of these AD risk genes were obtained from studies using the hippocampus. The gene expression patterns of the hippocampus generally differ from those of the prefrontal cortex examined in this study. Thus, the results obtained in this study were quite different from those of previous studies.

(a) “Table_S6a.xlsx”

(b) “Table_S6b.xlsx”

**Table S7**. Significant DEGs among patients with NCI, MCI, and AD.

“Table_S7.xlsx”

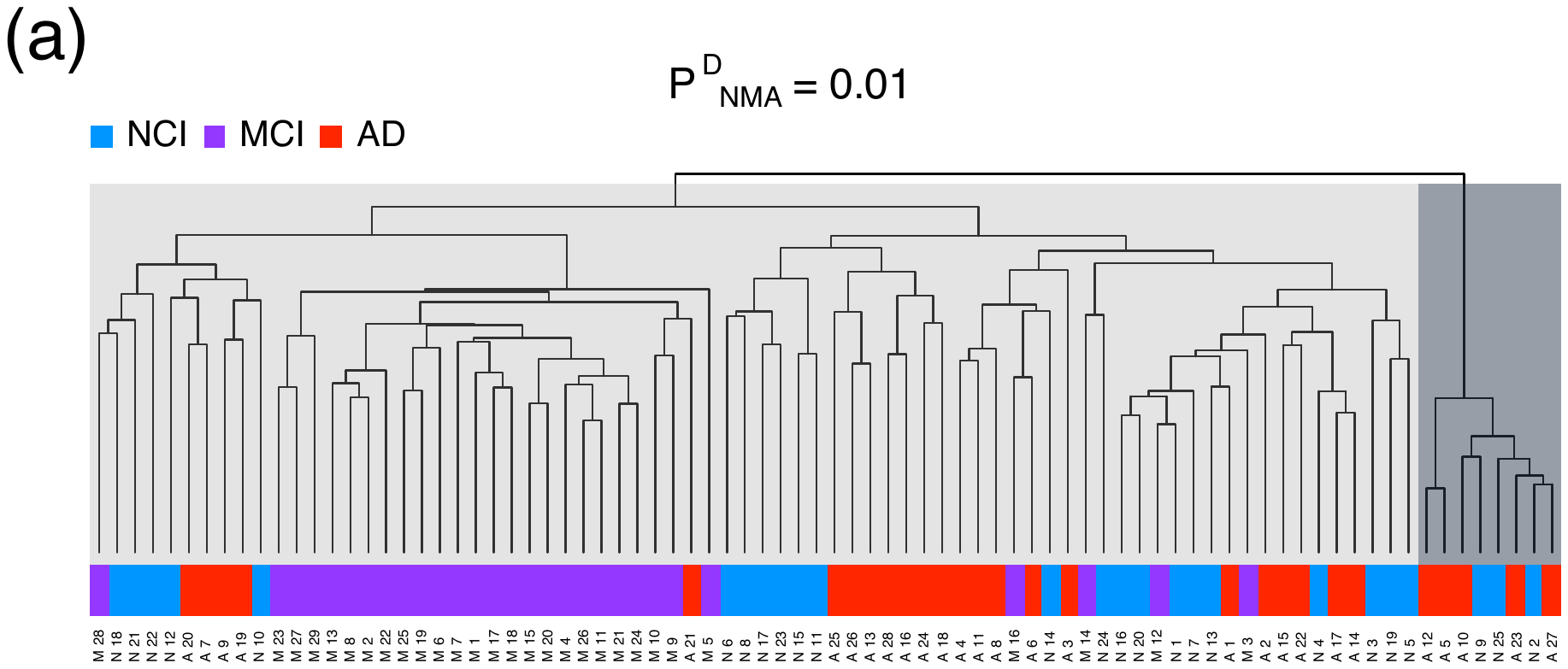

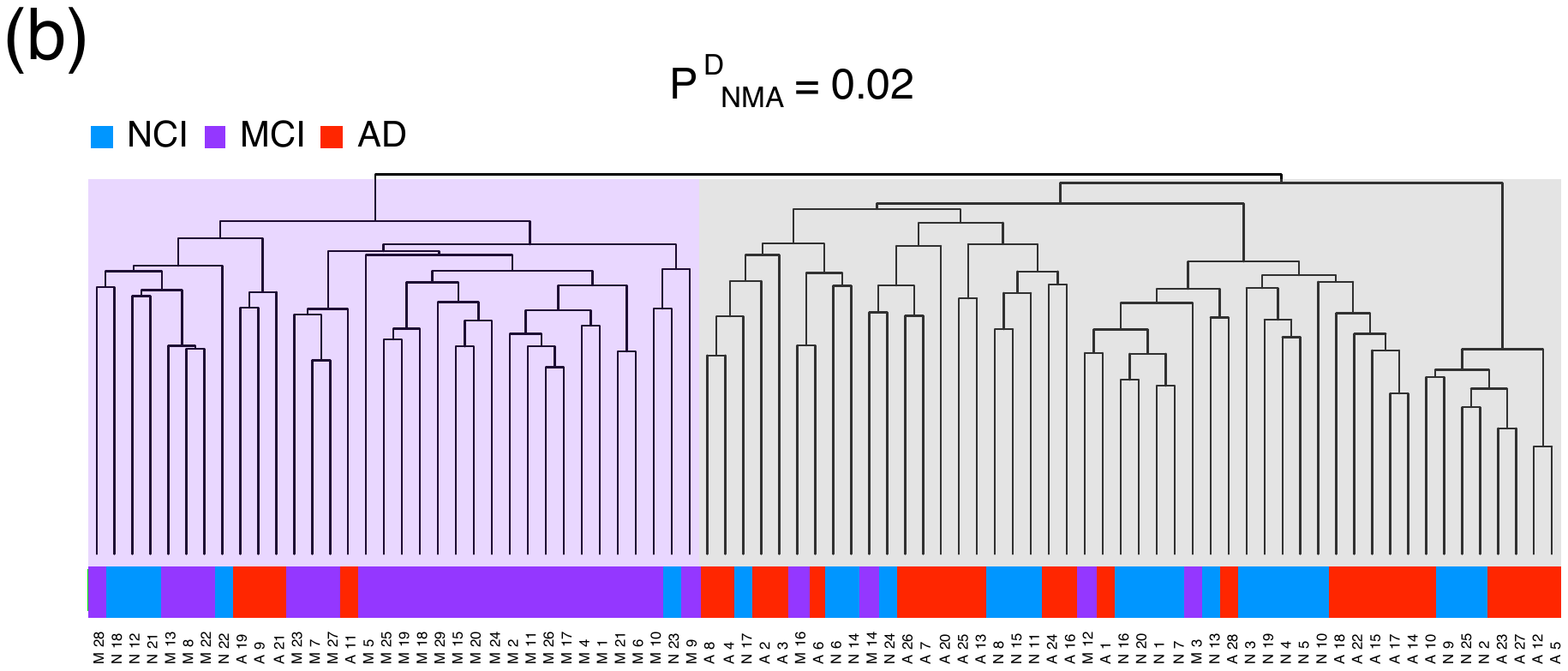

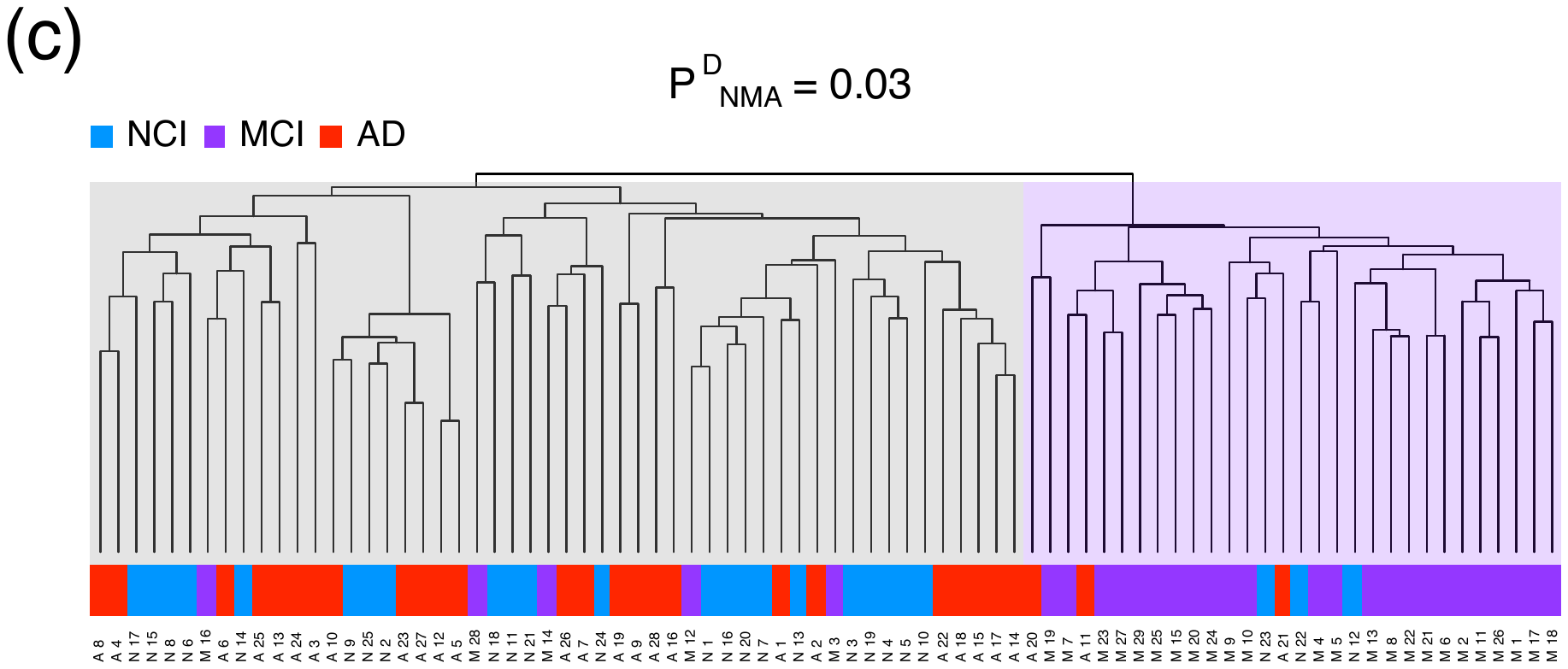

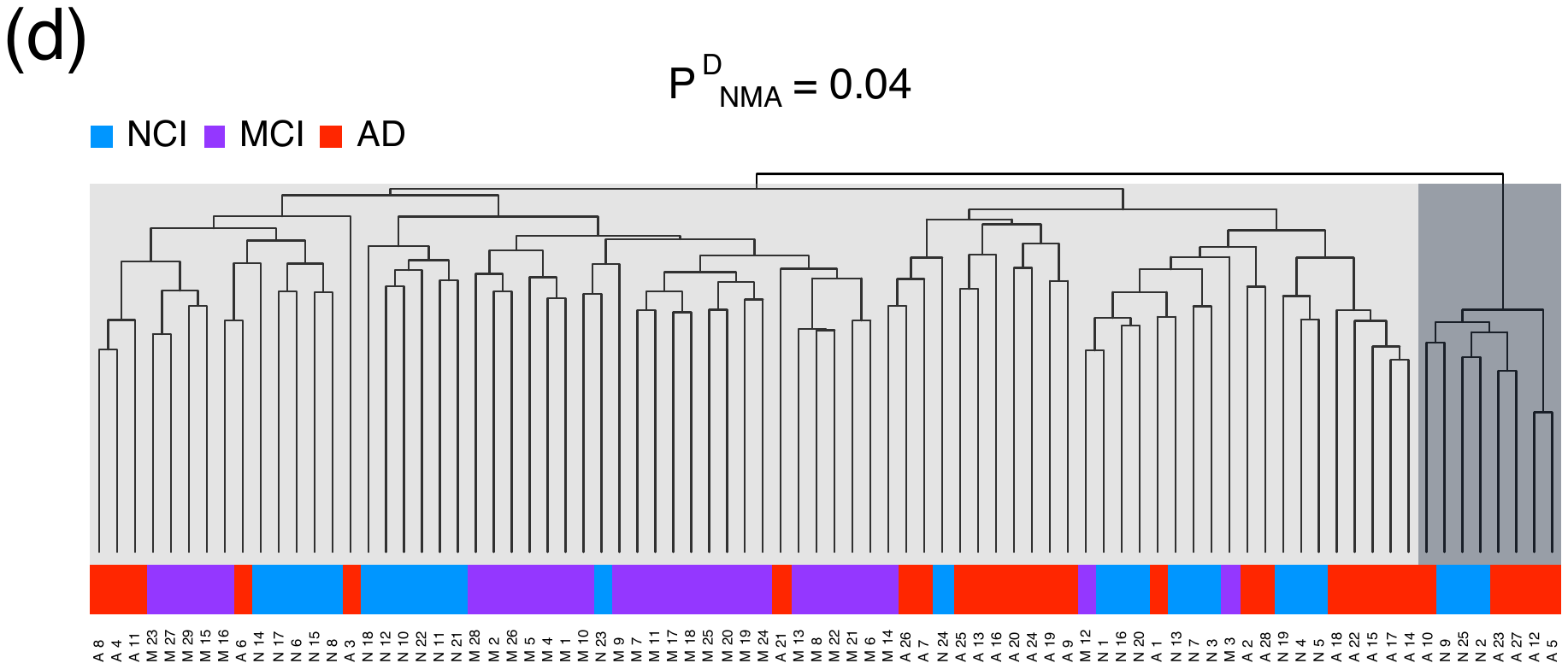

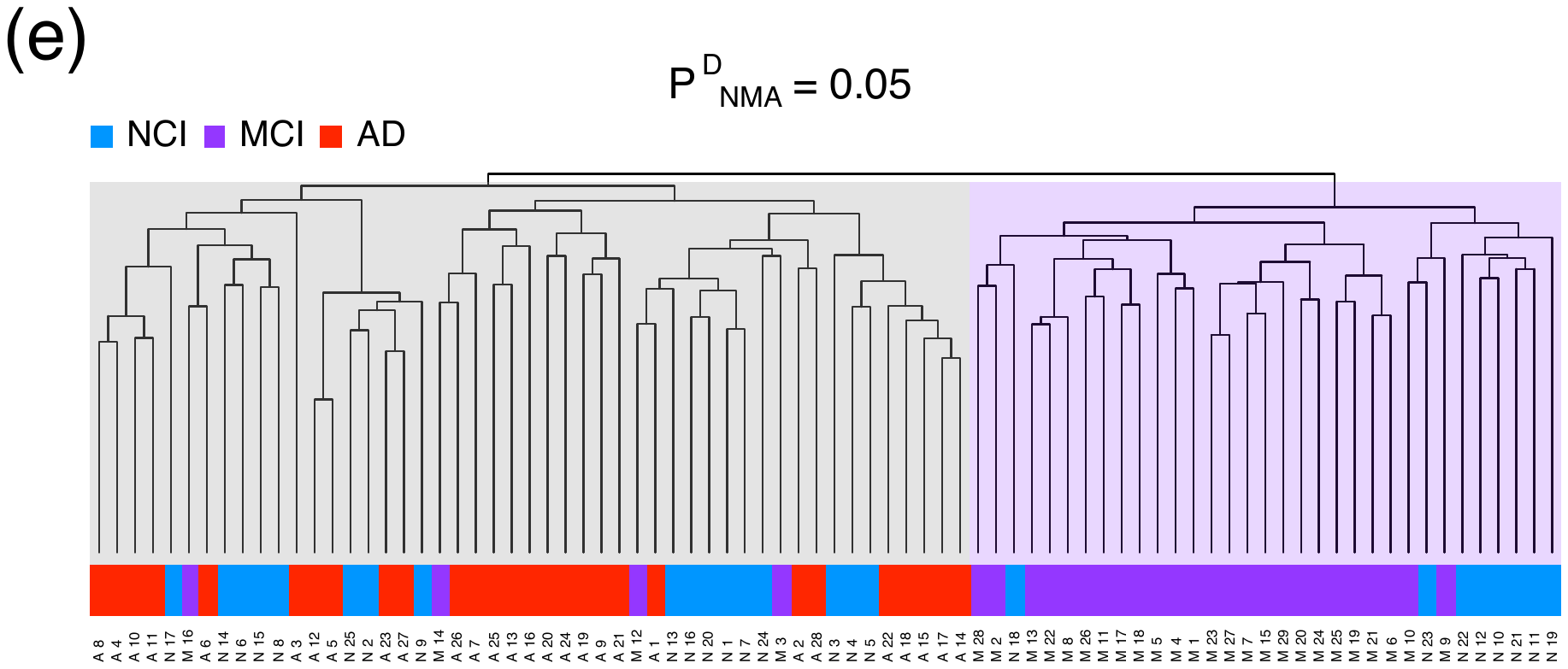

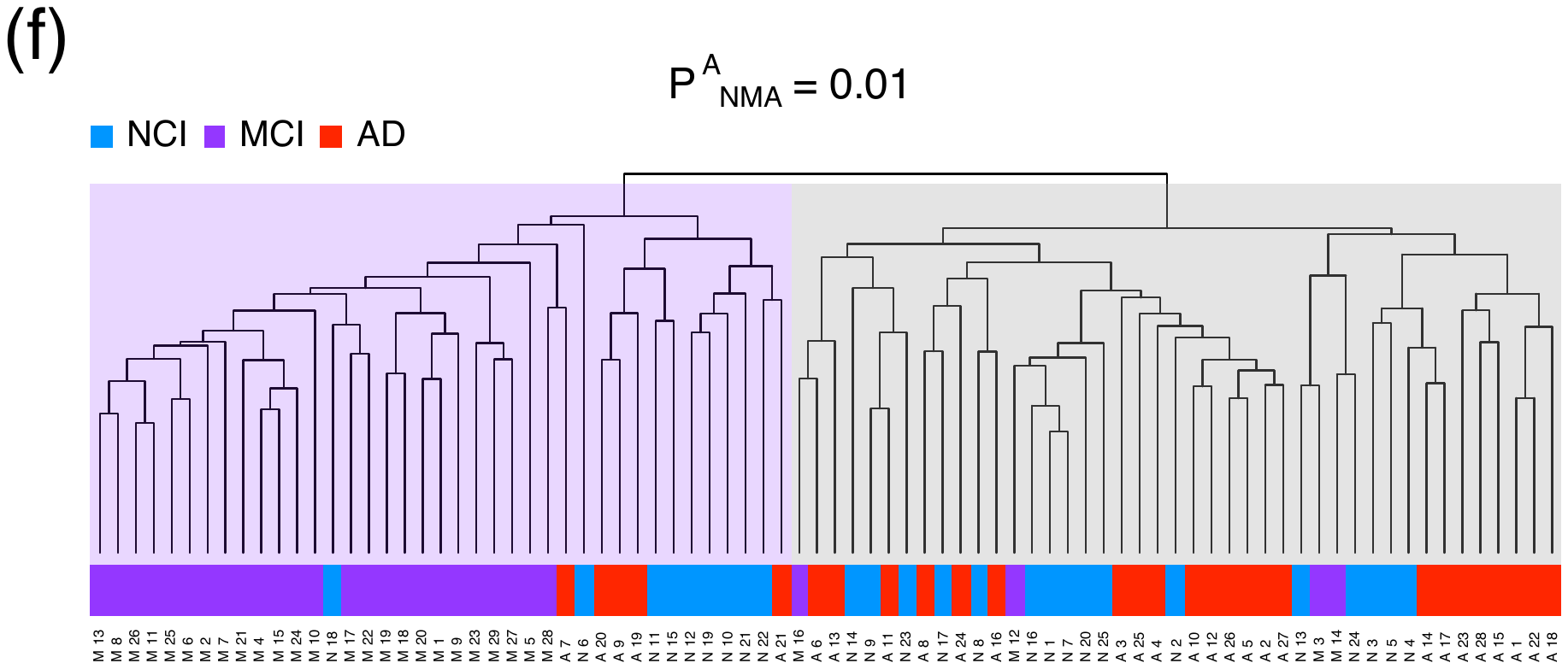

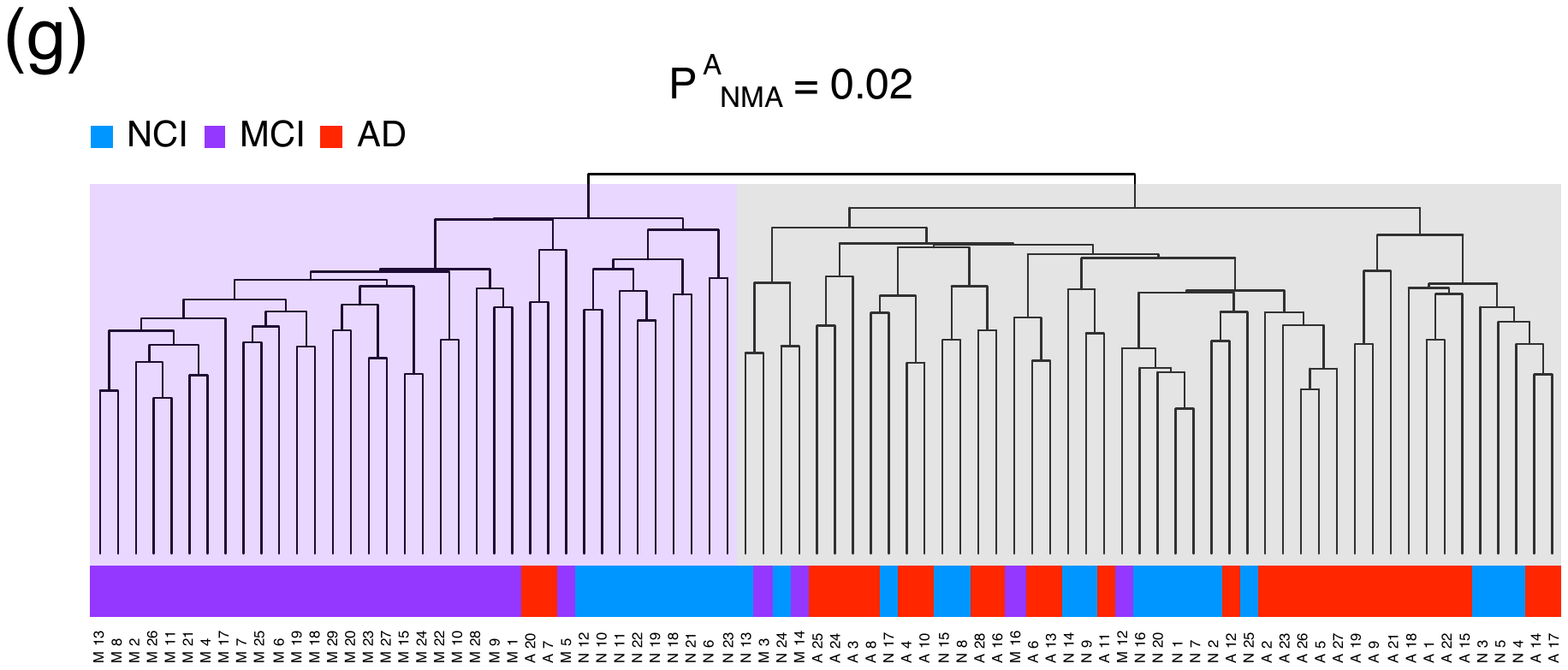

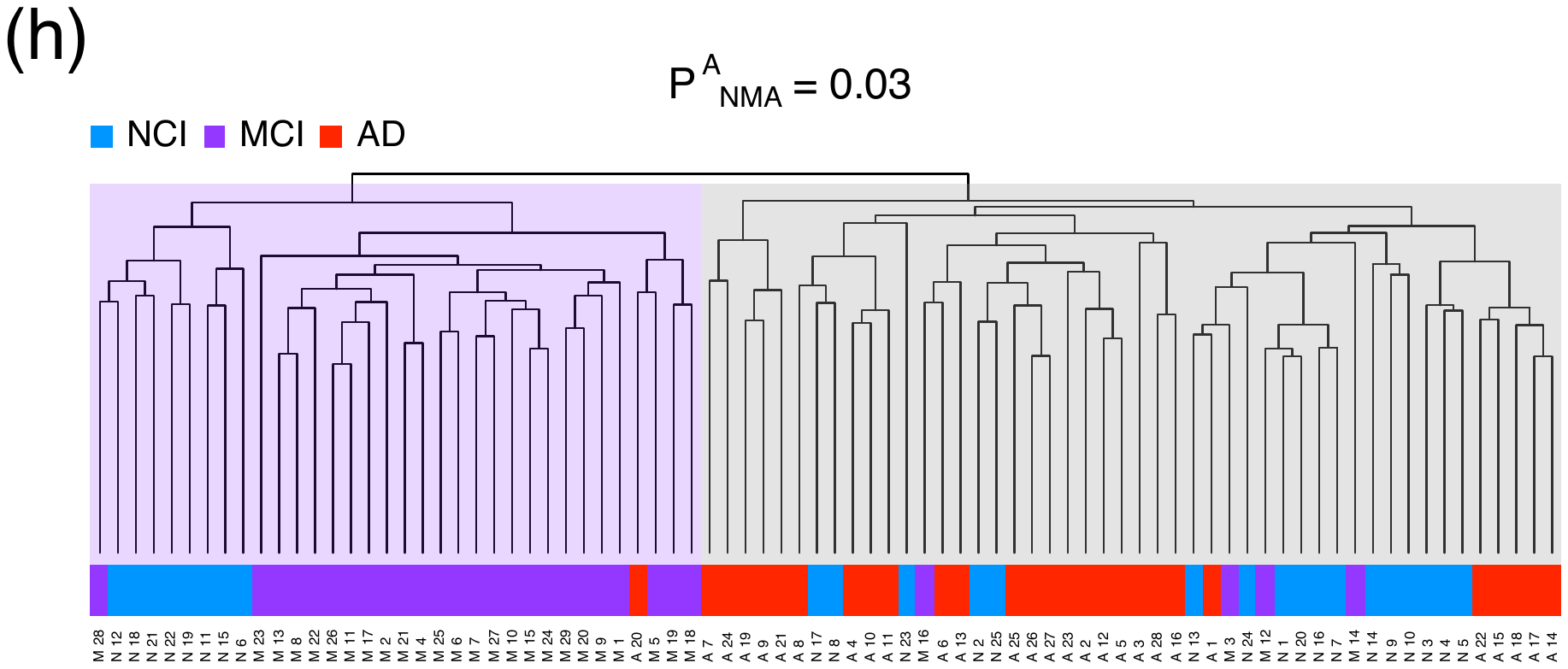

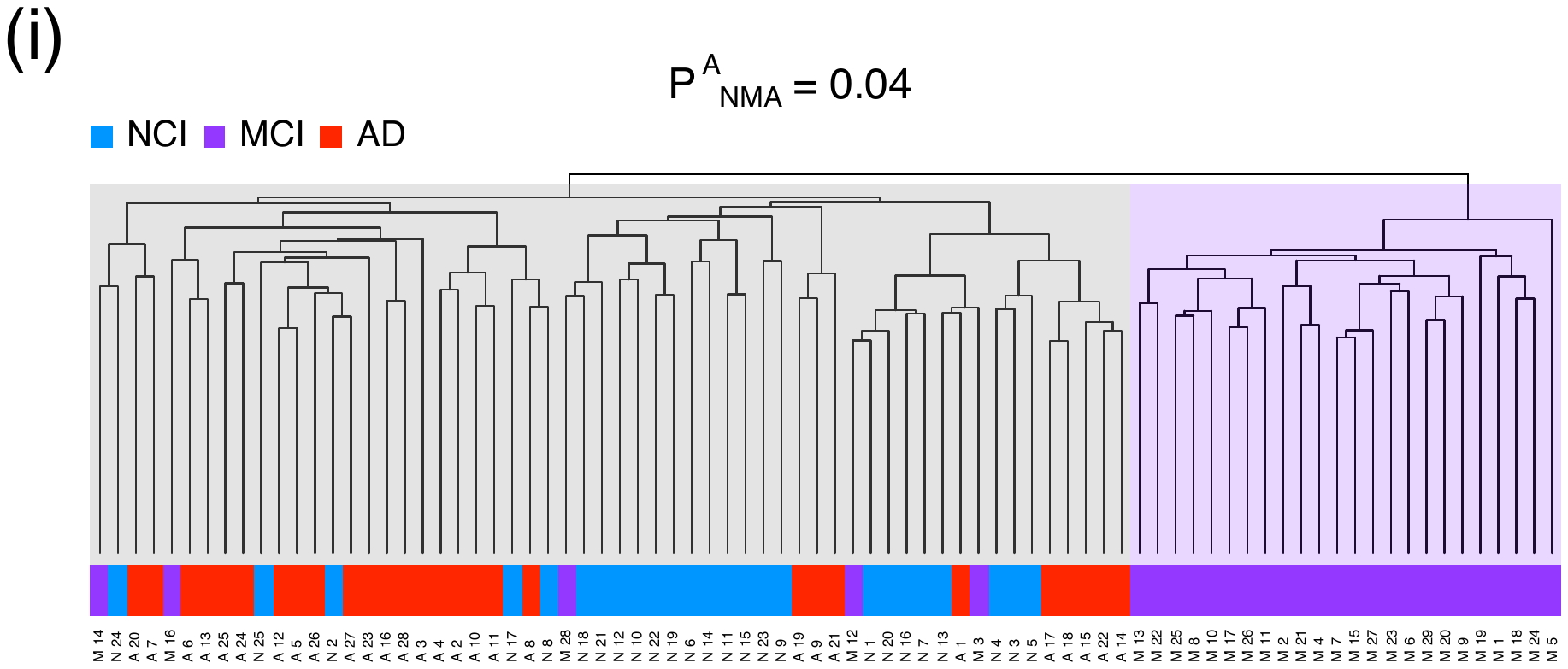

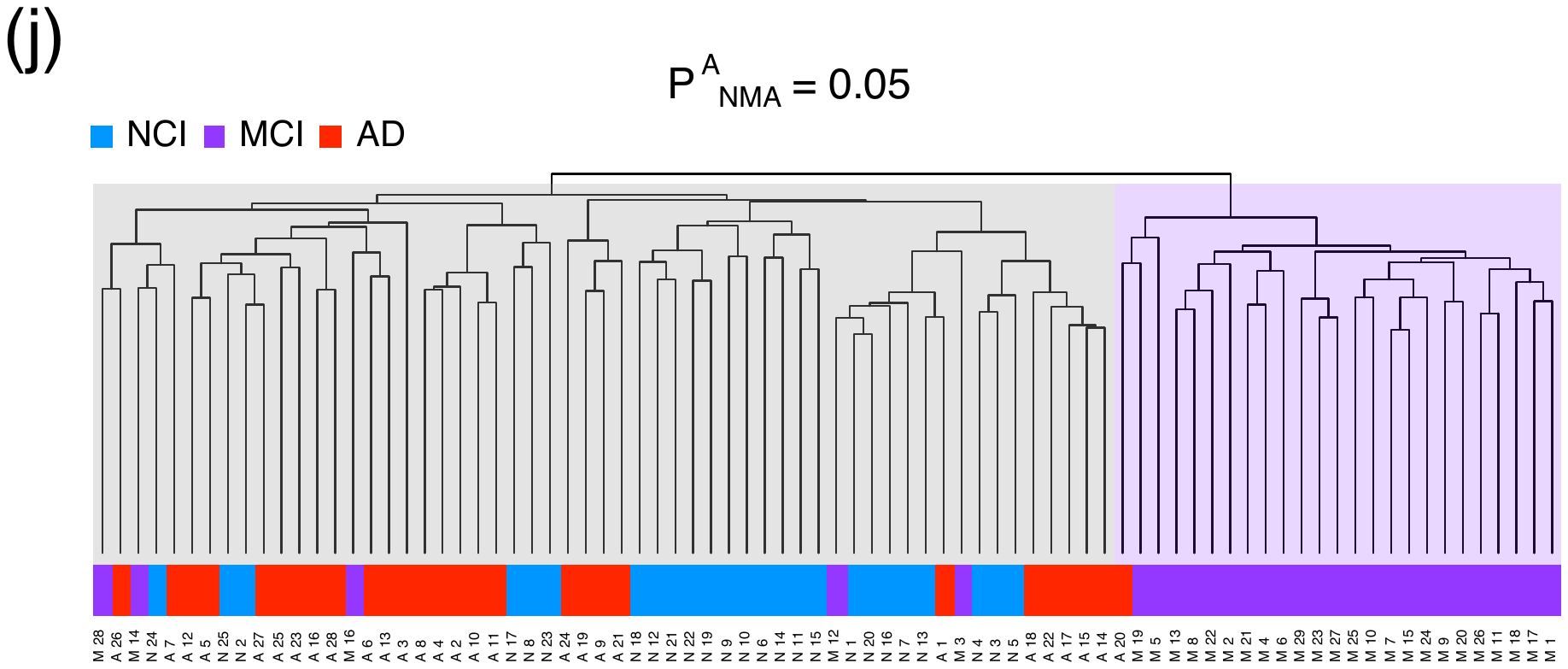

**Fig. S1 Dendrograms of the hierarchical clustering among the investigated individuals.** Dendrograms of the hierarchical clustering among individuals when P^D^_NMA_ = 0.01 (a), 0.02 (b), 0.03 (c), 0.04 (d), and 0.05 (e), and when P^A^_NMA_ = 0.01 (f), 0.02 (g), 0.03 (h), 0.04 (i), and 0.05 (j).

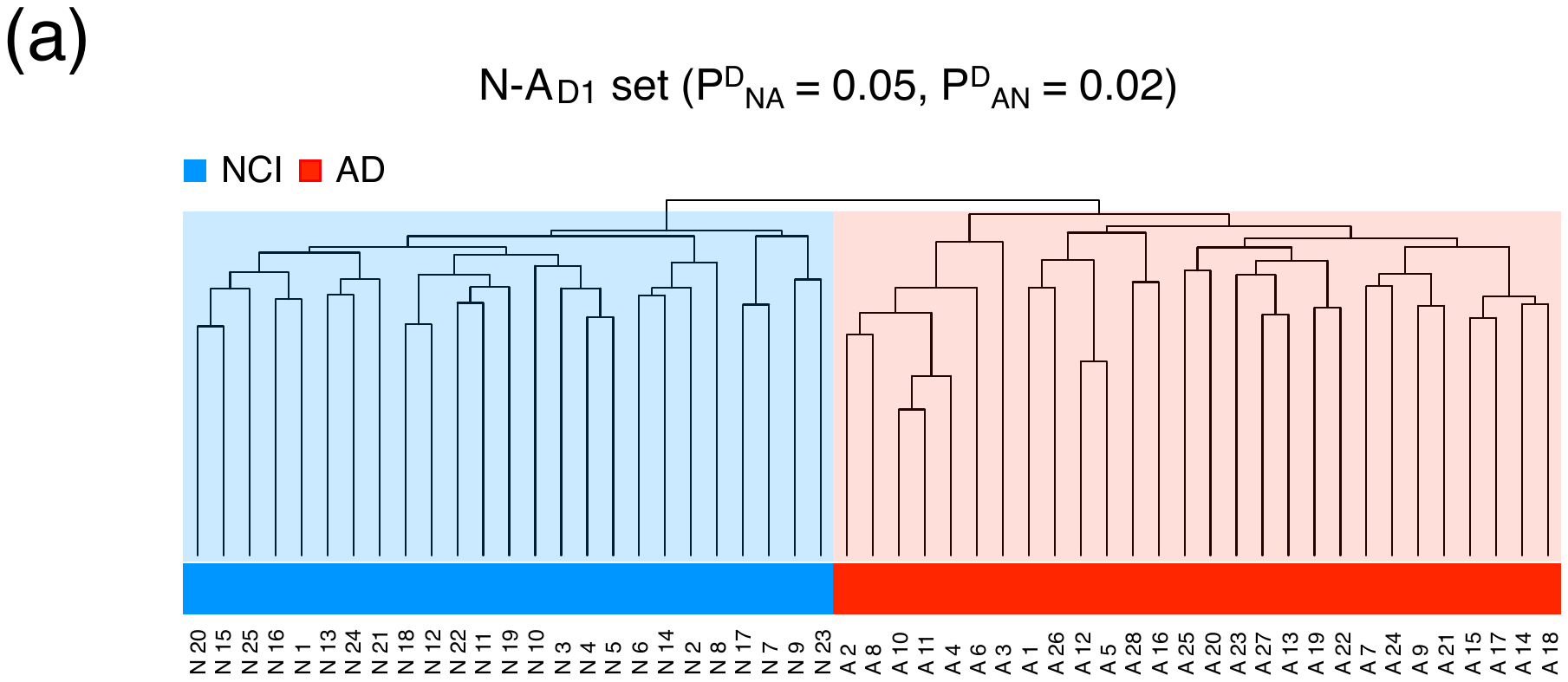

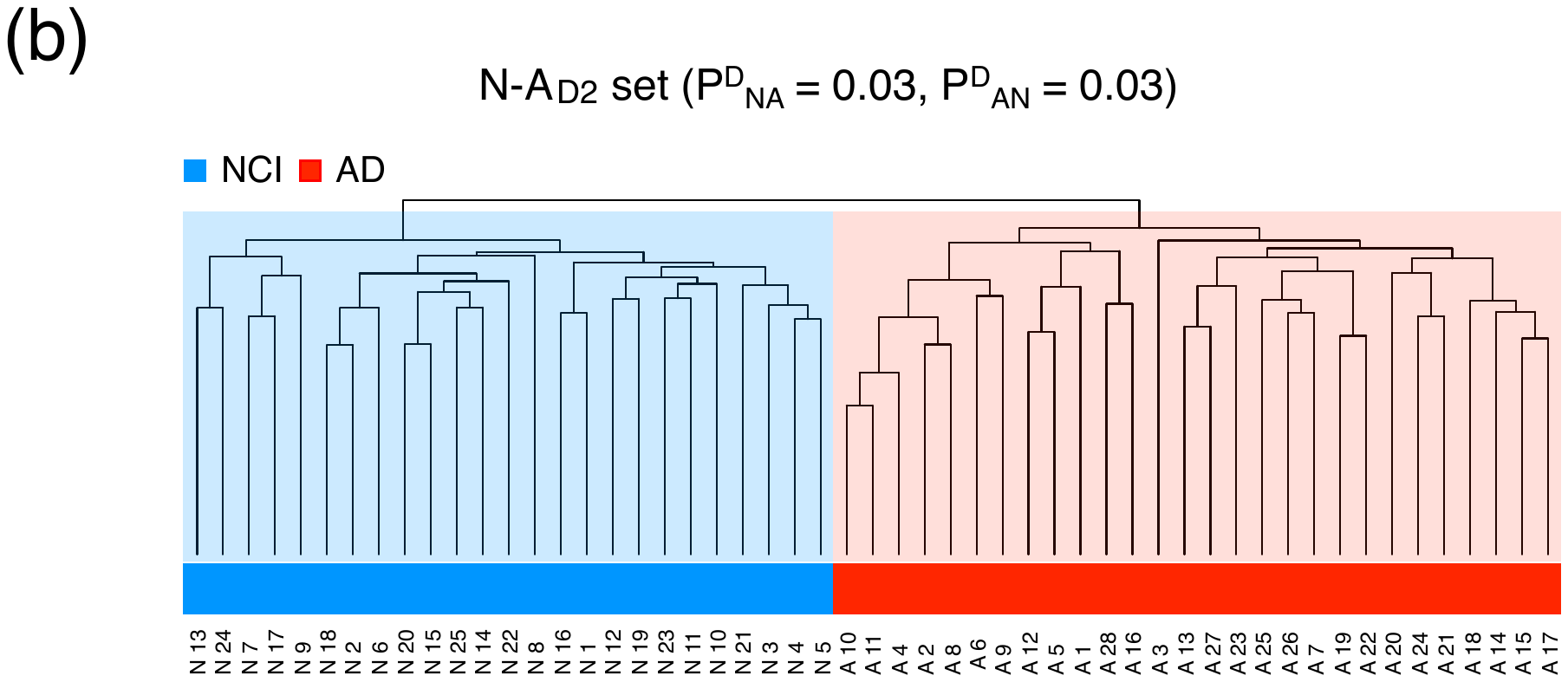

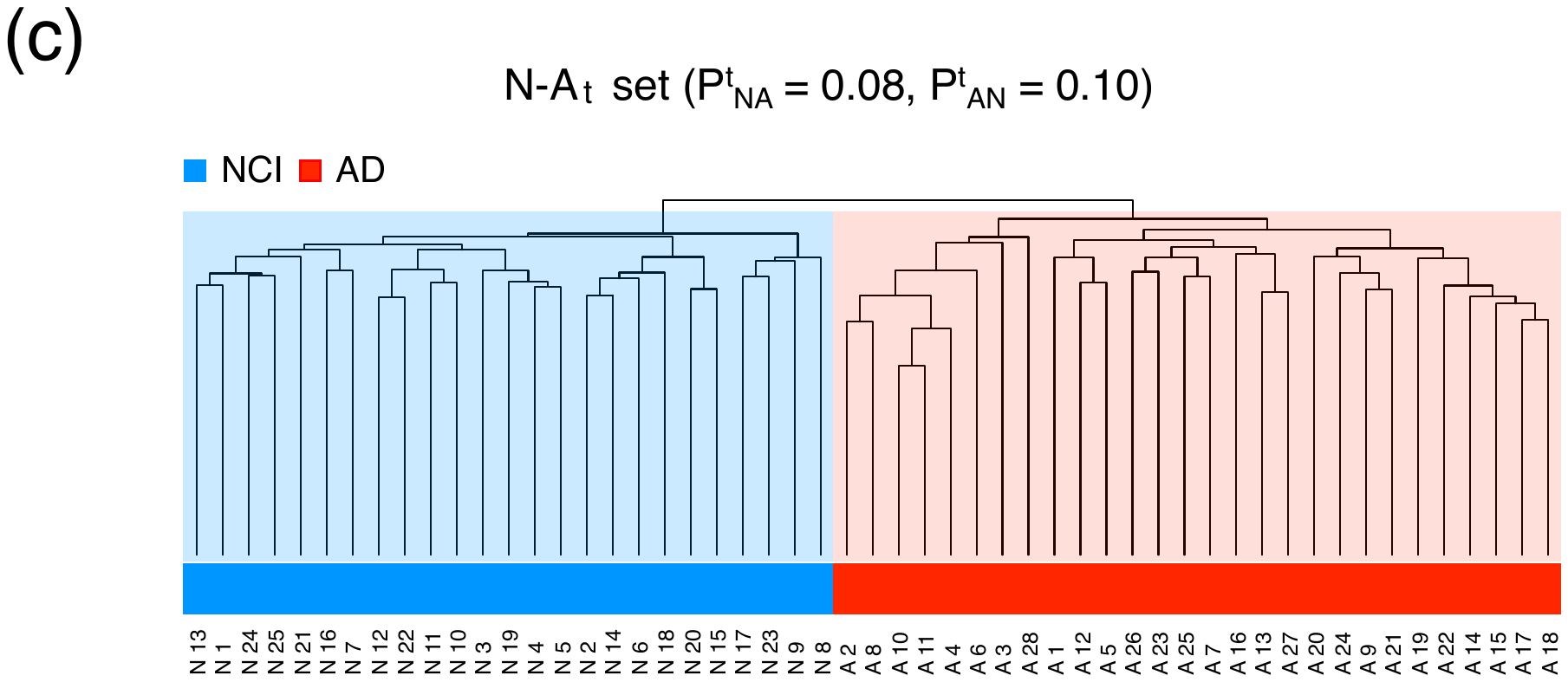

**Fig. S2** Dendrograms of the hierarchical clustering among patients with NCI and AD obtained using the N-A_D1_ (a), N-A_D2_ (b), and N-A_t_ (c) sets.

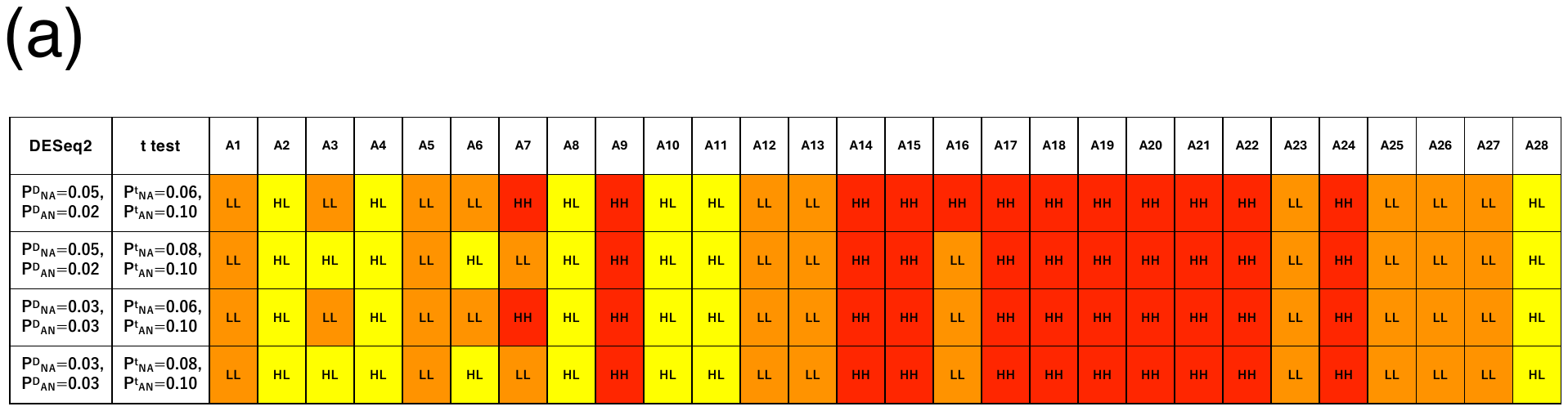

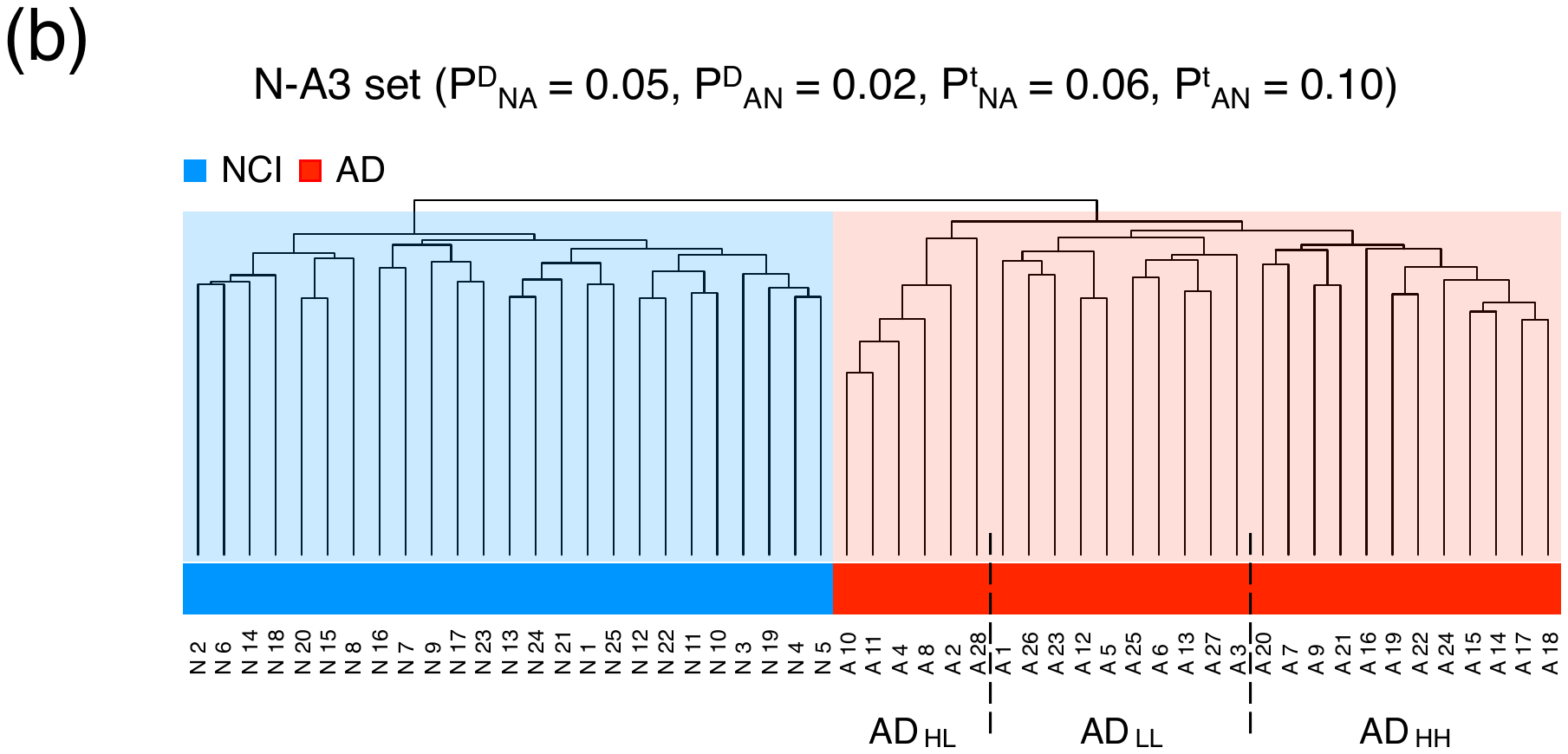

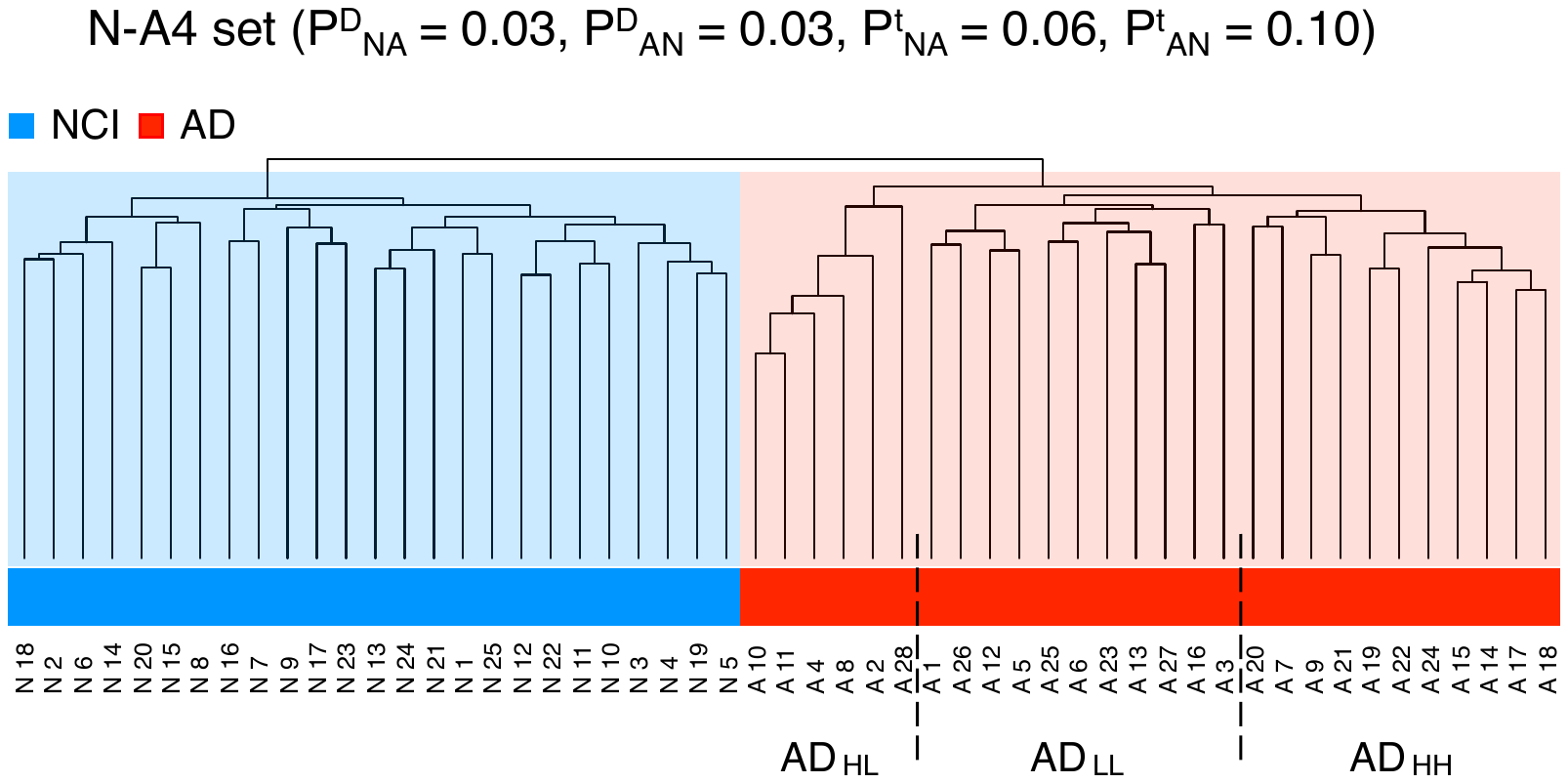

**Fig. S3.** Dependencies of belonging to each subgroup of patients with AD based on the P^D^_NA_, P^D^_AN_, P^t^_NA,_ and P^t^_AN_ values (a). Dendrograms of the hierarchical clustering among patients with NCI and AD obtained using the N-A3 (b, upper panel) and N-A4 (b, lower panel) sets.

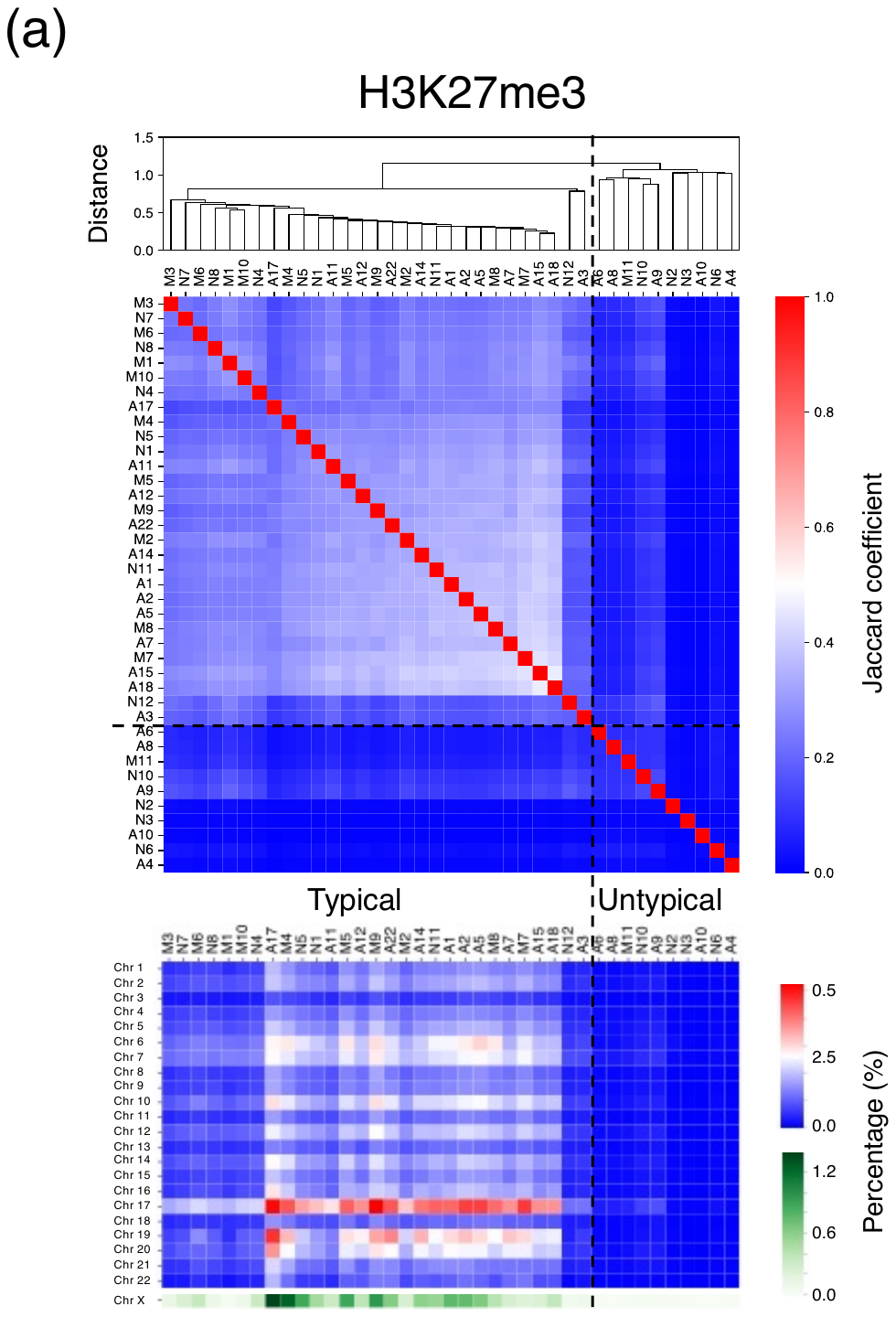

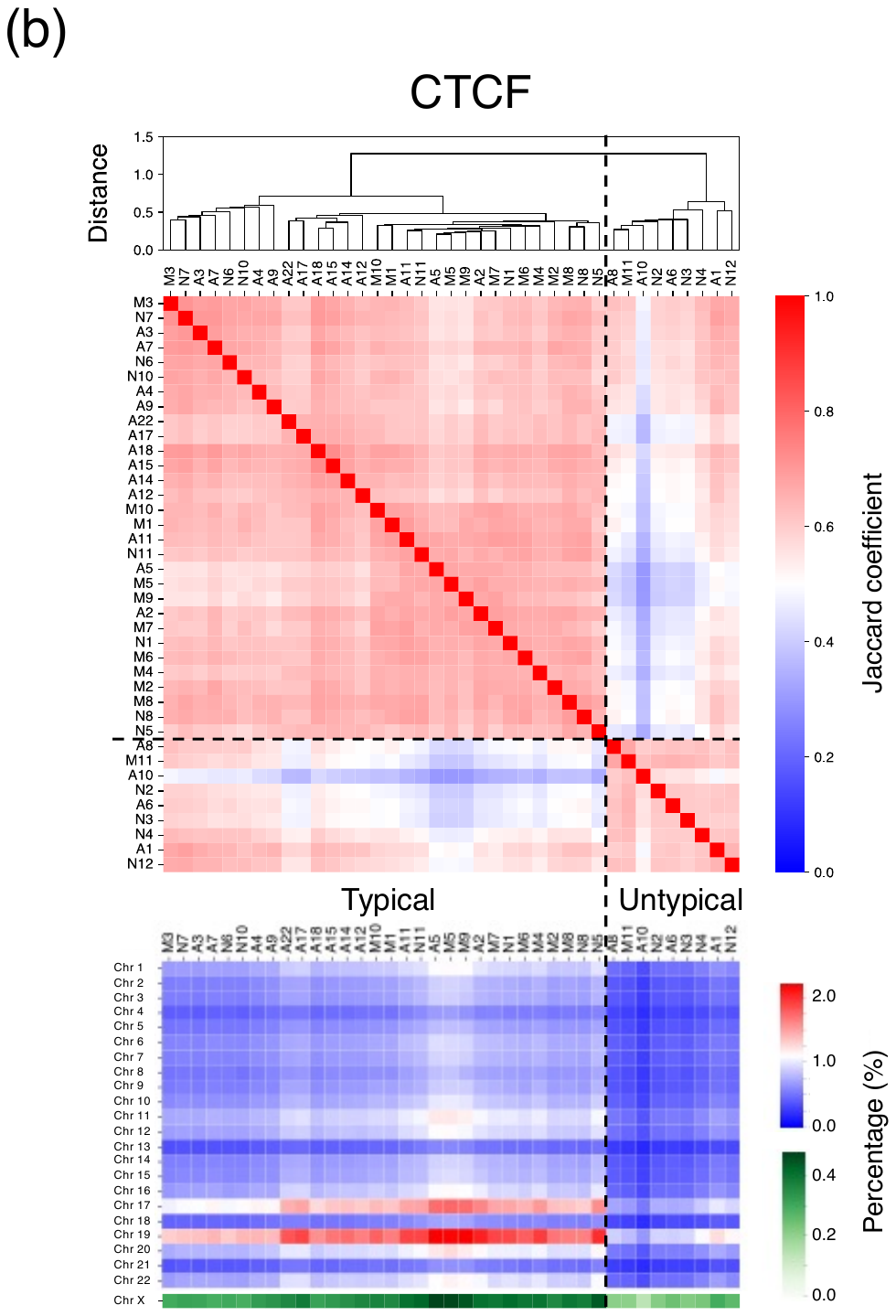

**Fig. S4** Jaccard coefficient of the genome-wide distributions of H3K27me3 (a), and CTCF (b) among individuals. Individuals were divided into two groups at the first divergence of the dendrogram. The epigenetic state exhibited by individuals belonging to the group containing the larger number of individuals was regarded as the "typical" epigenetic state, whereas that exhibited by individuals belonging to the group containing the fewer individuals was regarded as the "untypical" epigenetic state. Dashed lines indicate the boundaries between groups.

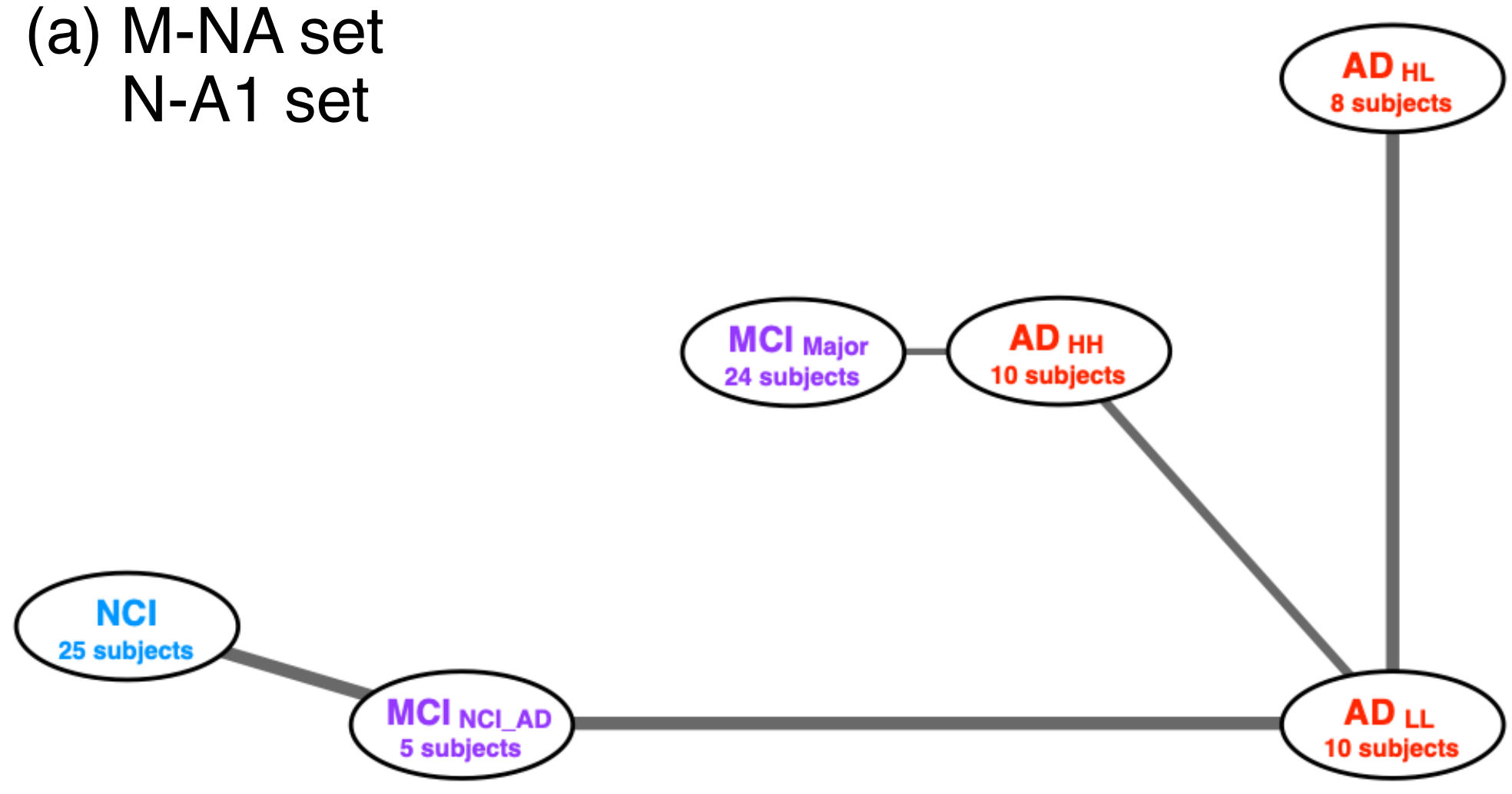

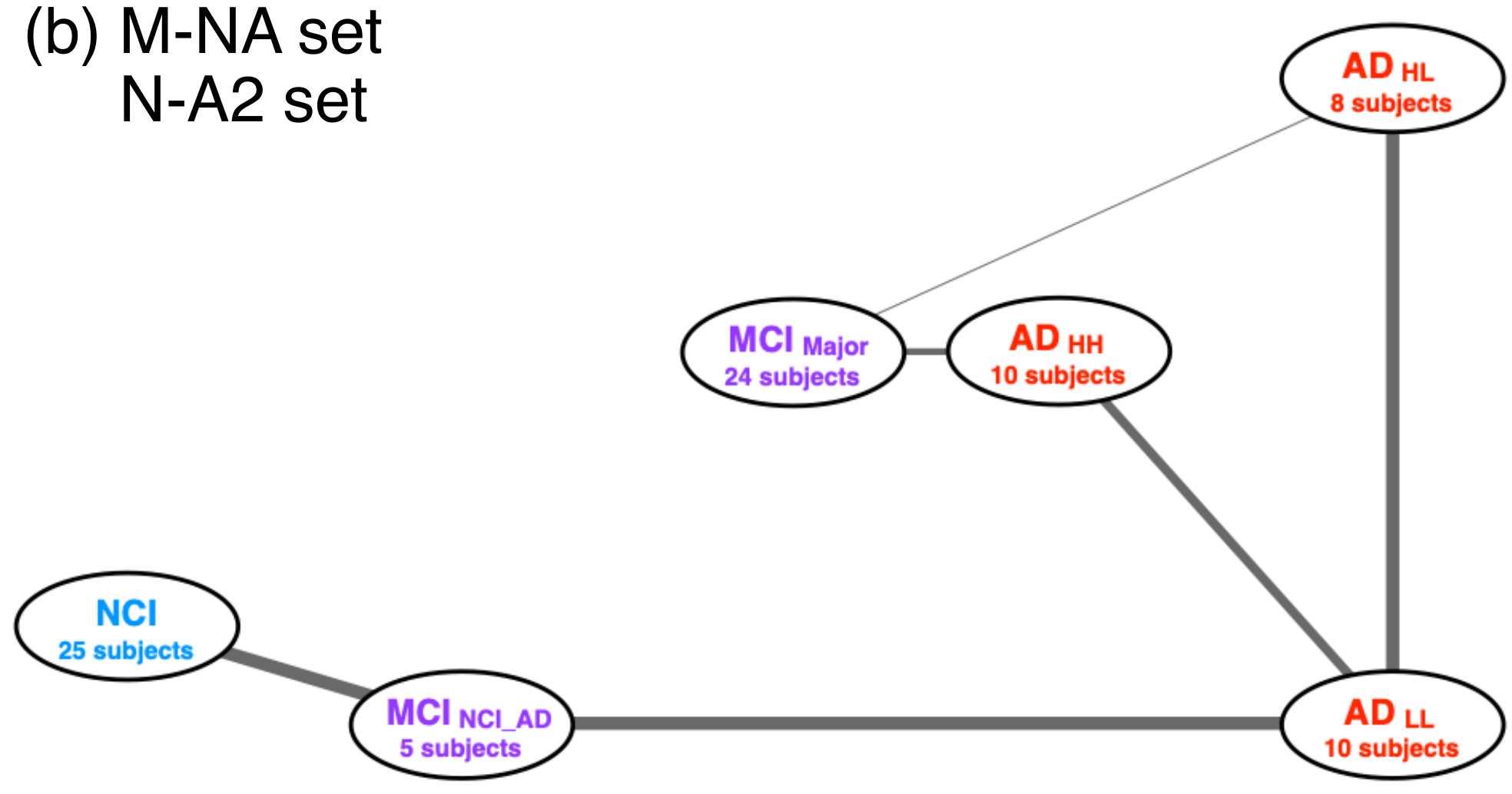

**Fig. S5** Adjacency network obtained using the M-NA and N-A1 sets (a) and that constructed using the M-NA and N-A2 sets (b).

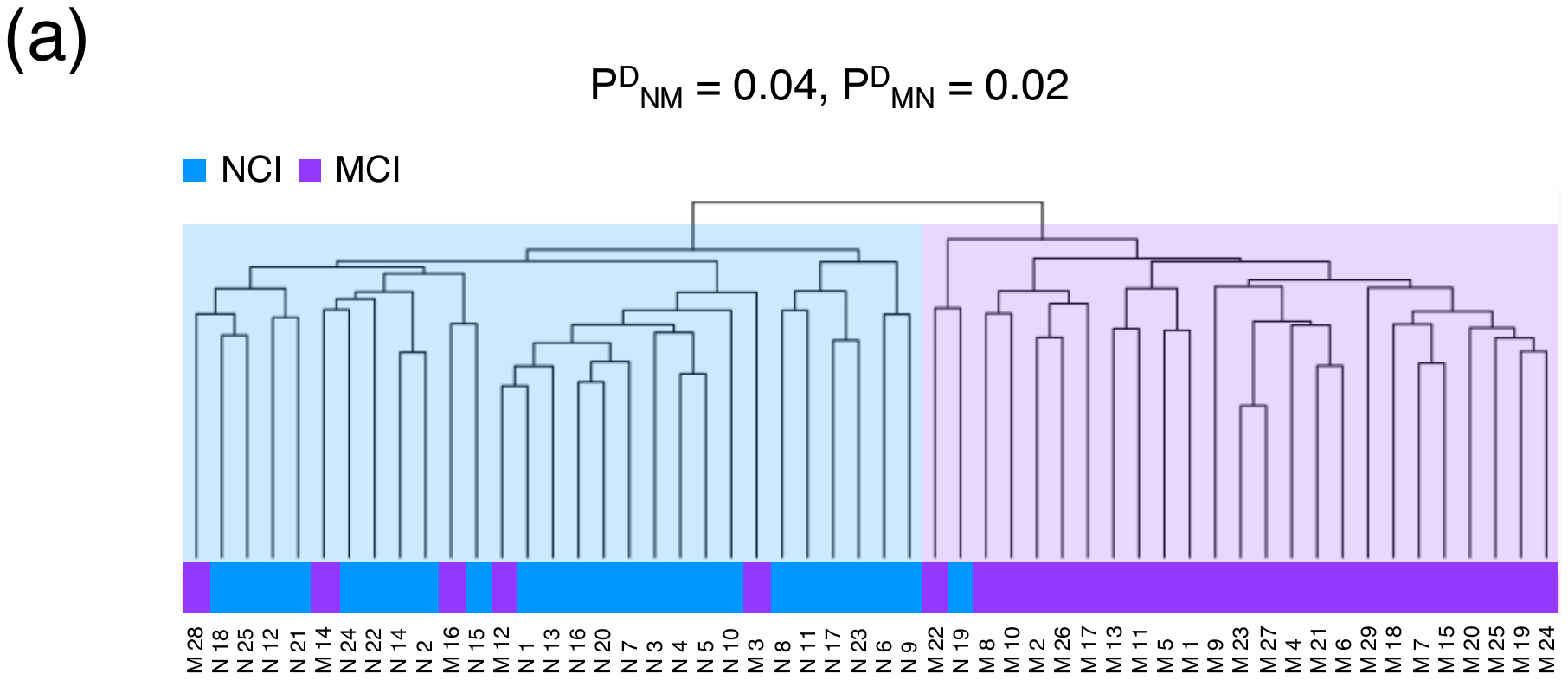

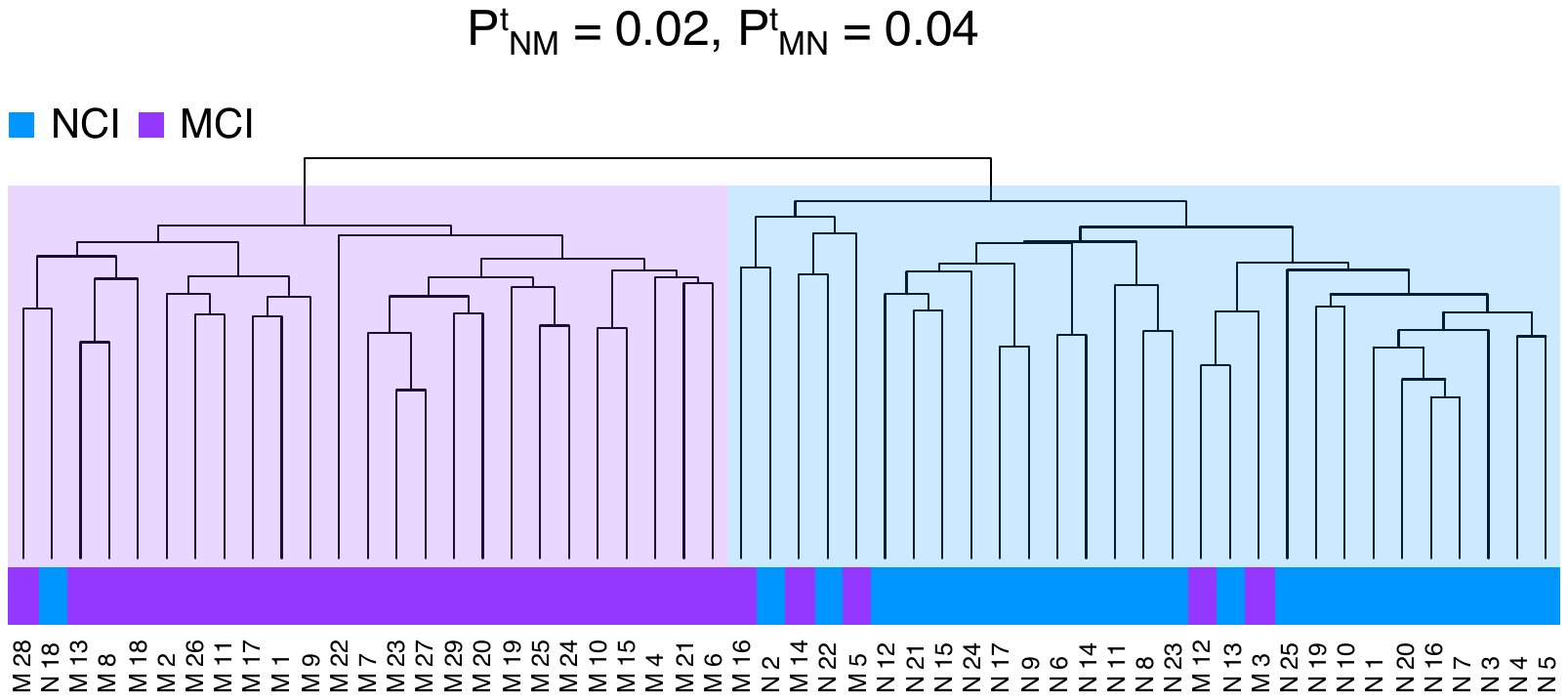

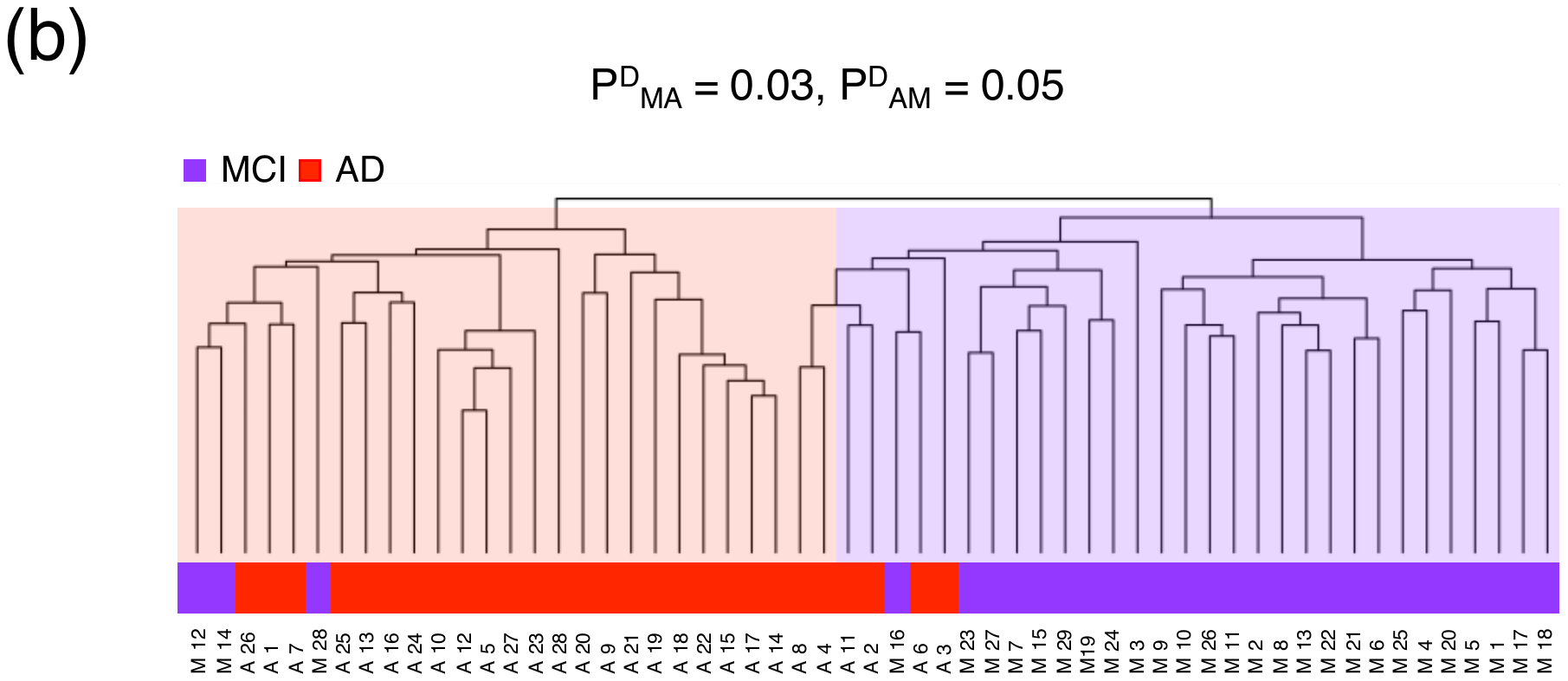

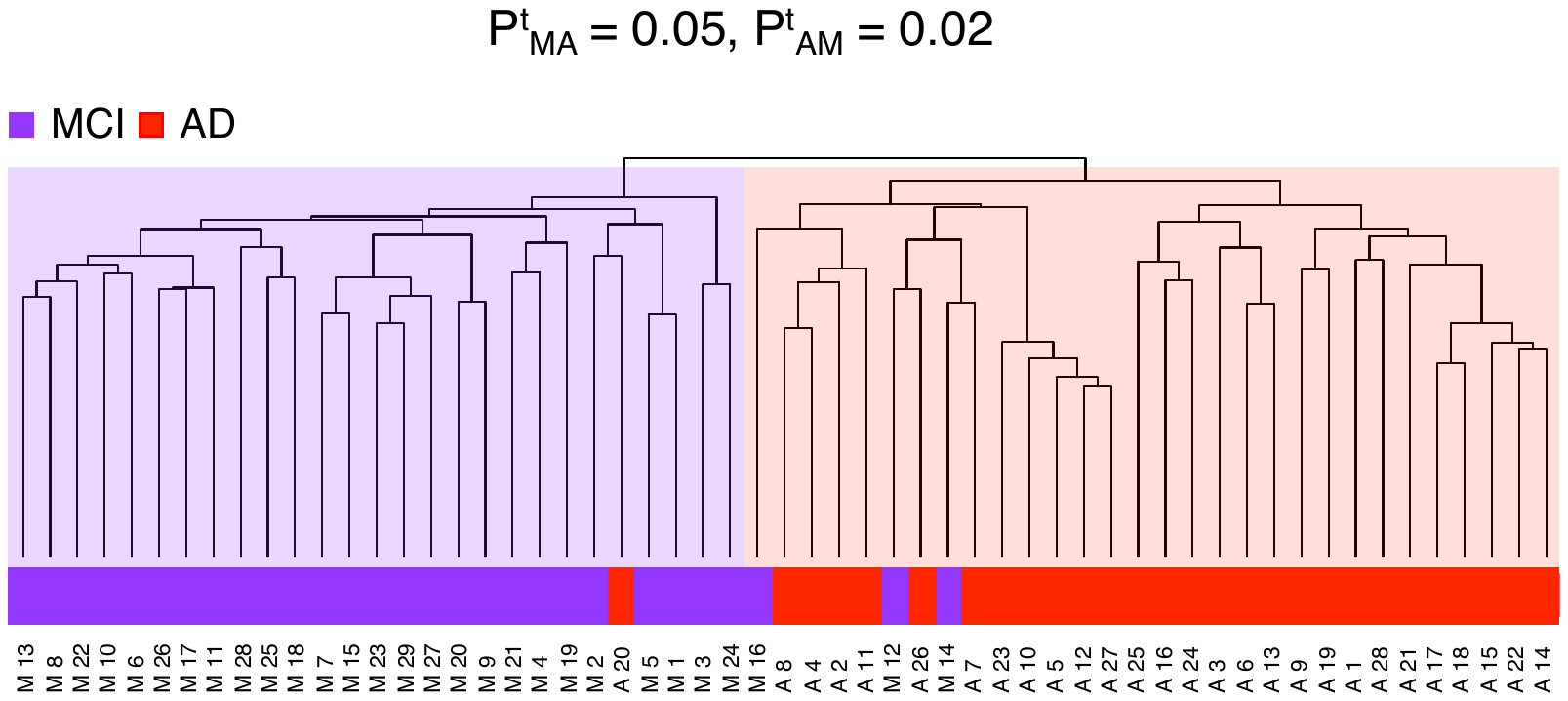

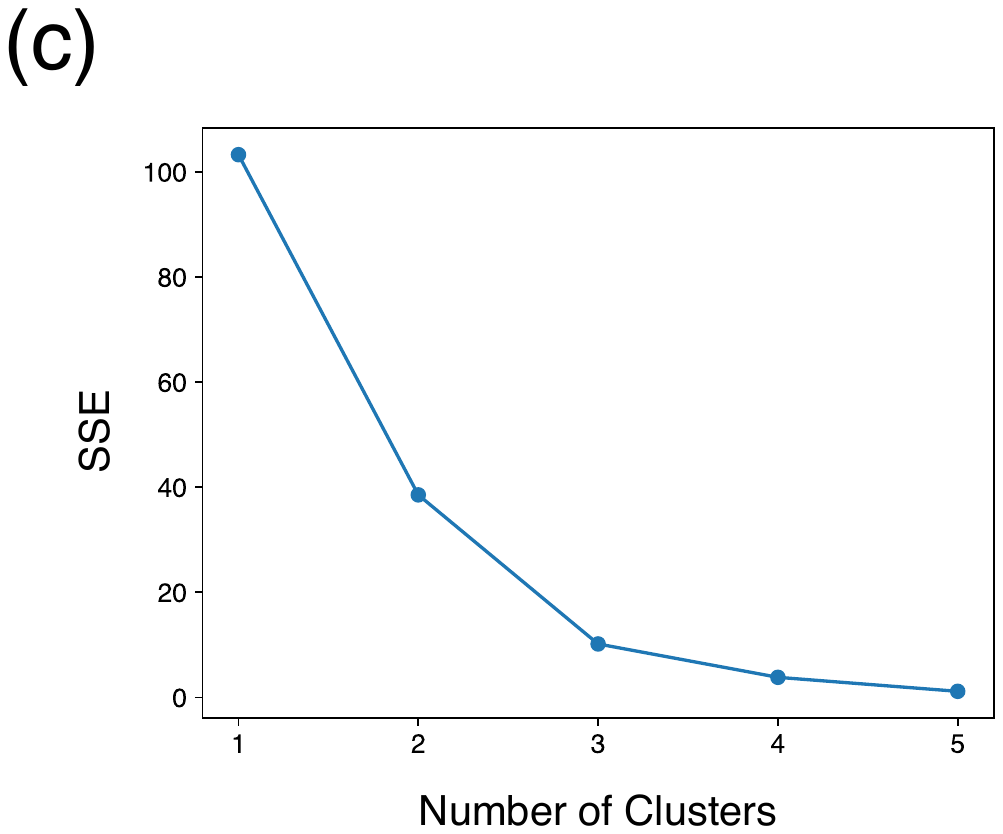

**Fig. S6** Examples of the hierarchal clustering of patients with NCI and MCI (a) and patients with MCI and AD (b). The sum of squared errors (SSE) as a function of the number of clusters for the k-means clustering of MCI_Major_ individuals using the number of times each individual belonged to the NCI or AD clusters (c). This plot shows the inferred number of clusters (= 3).

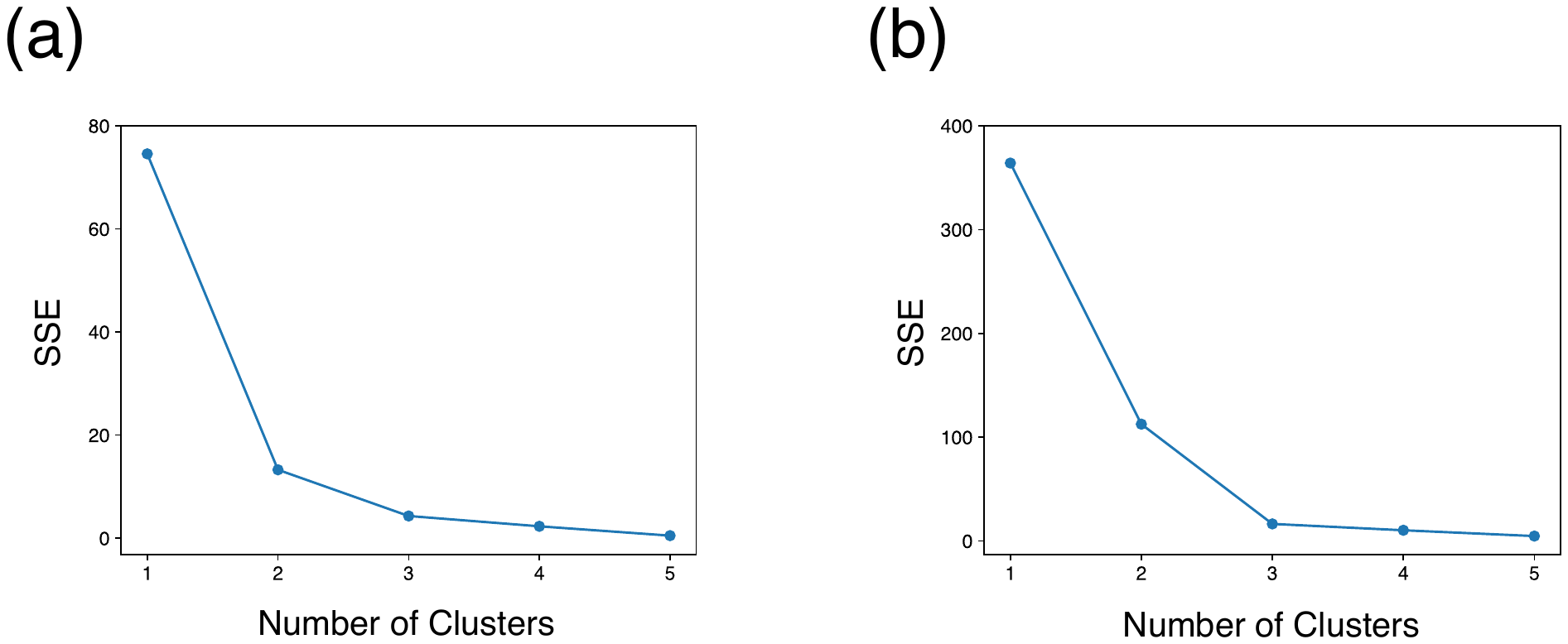

**Fig. S7** The sum of squared errors (SSE) as a function of the number of clusters for k-means clustering of patients with NCI using the number of times each individual belonged to the MCI cluster (a), and that of patients with AD using the number of times each individual belonged to the MCI cluster (b). The number of clusters = 2 was inferred in (a), while the number of clusters = 3 was inferred in (b).

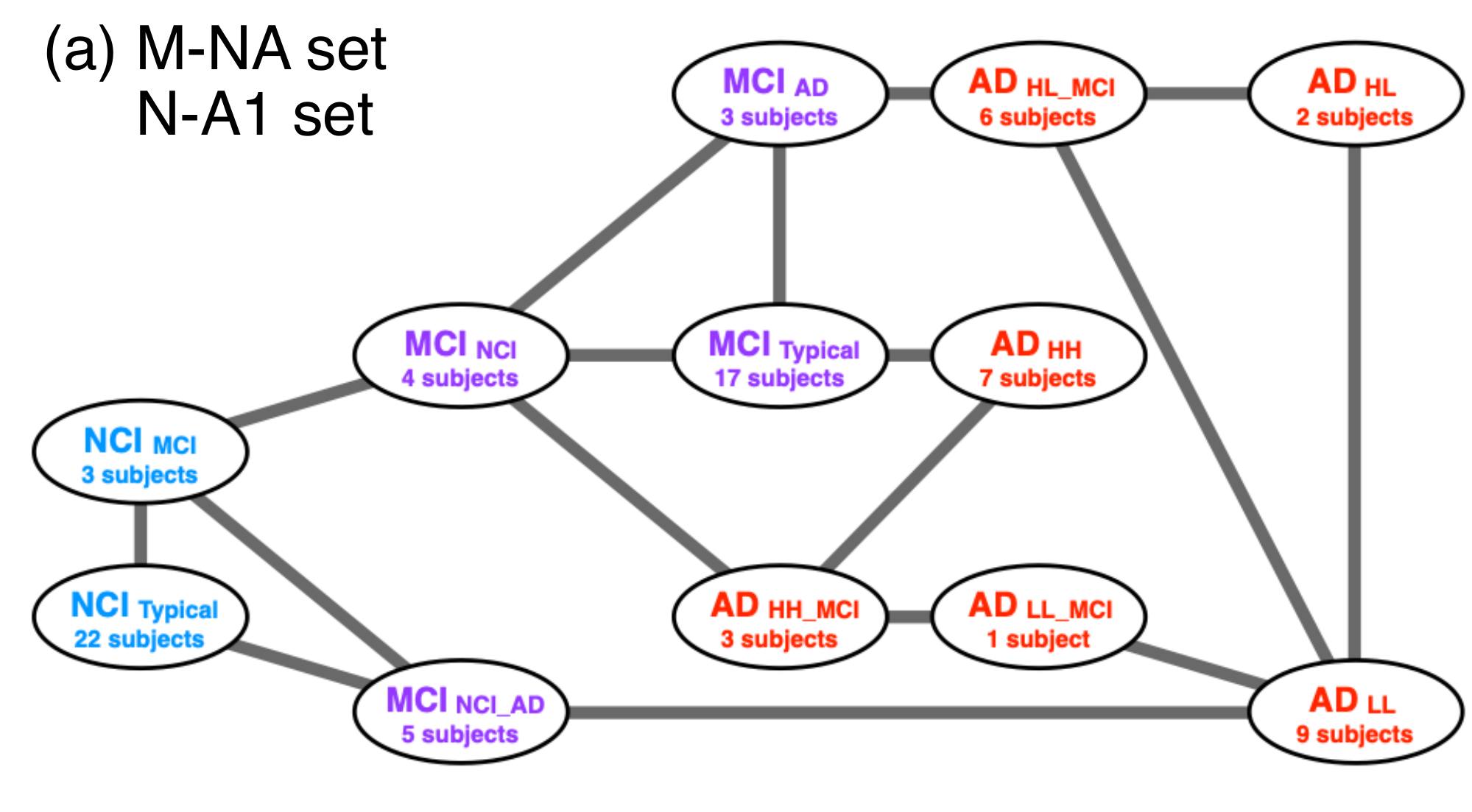

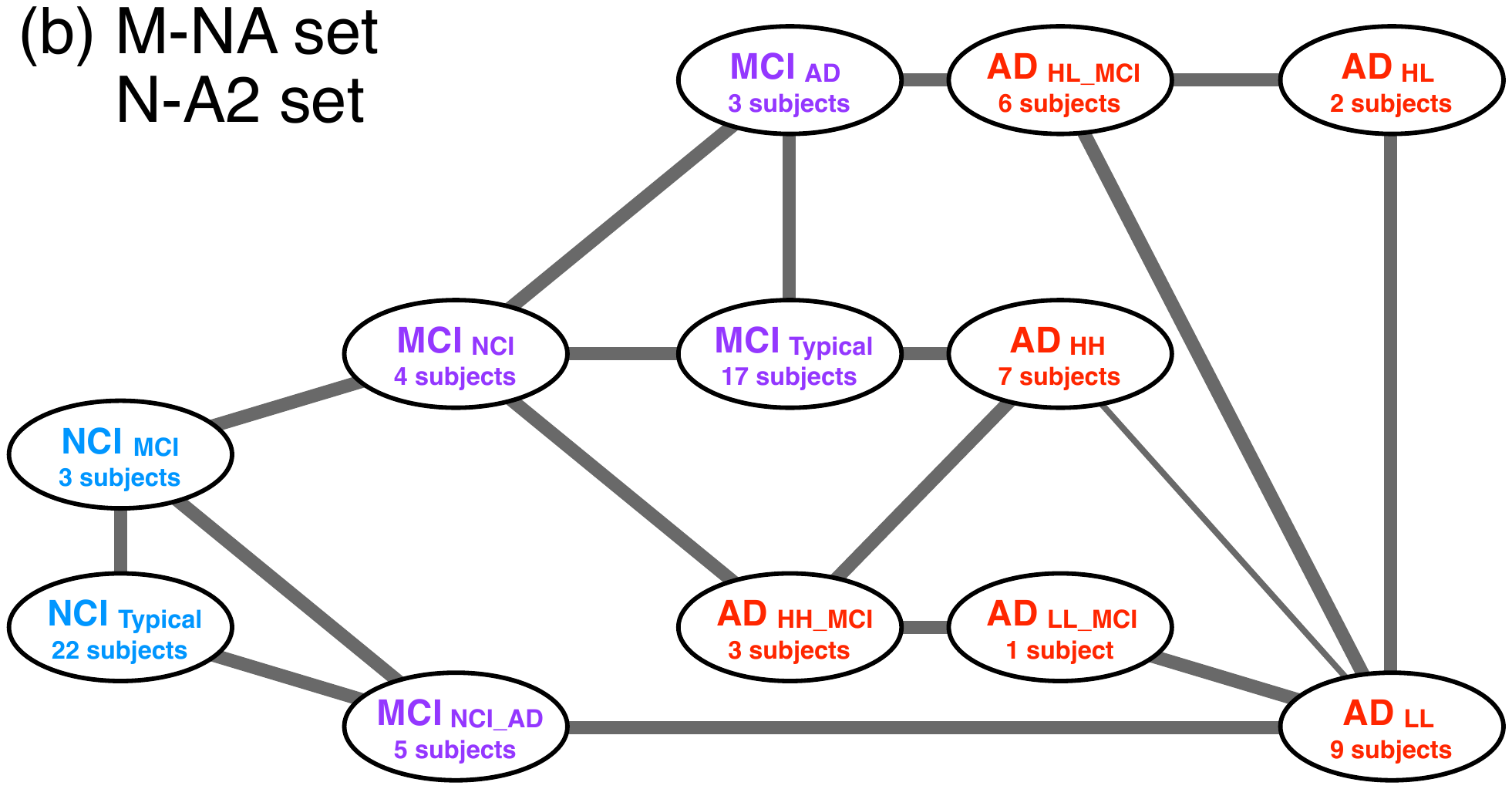

**Fig. S8** Adjacency network obtained using the M-NA and N-A1 sets (a) and that constructed using the M-NA and N-A2 sets (b).

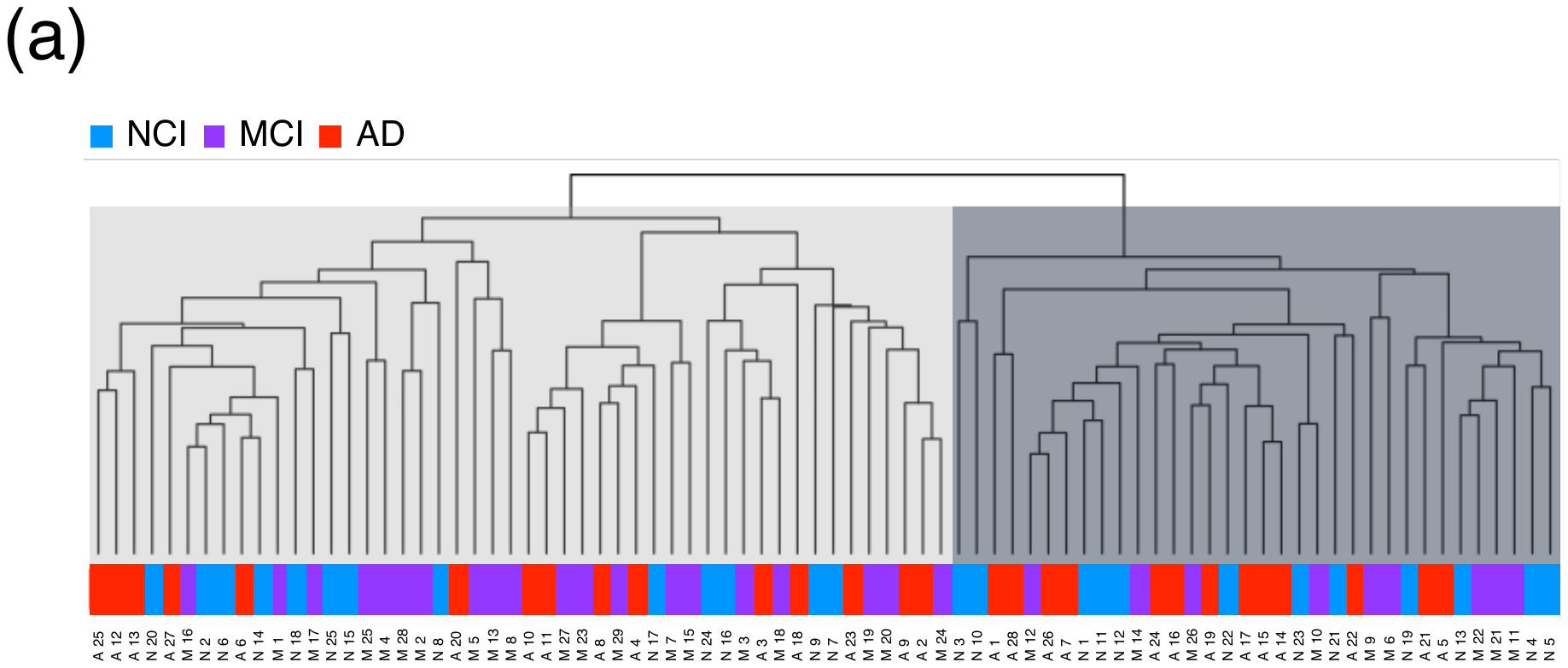

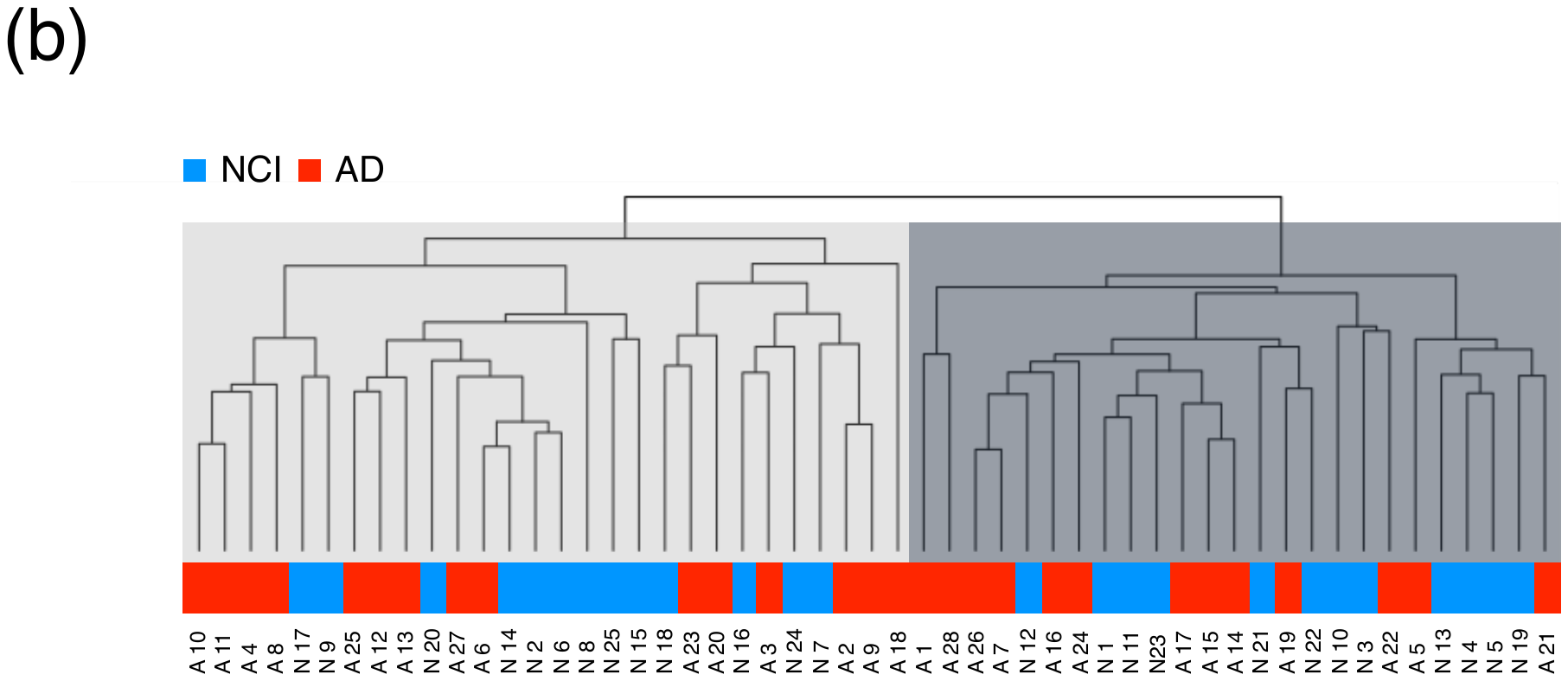

**Fig S9**. Results of the cluster analysis of patients with NCI, MCI and AD (a) and that of patients with NCI and AD (b) using AD risk genes previously reported in the literature (Hu et al. 2017; Xiang et al. 2018; Rahman et al. 2019; Yang et al. 2022). Many of these AD risk genes were obtained from studies using the hippocampus. The gene expression patterns of the hippocampus generally differs from that of the prefrontal cortex examined in this study. Thus, the results obtained in this study were quite different from those from previous studies.
